# Dose-dependent thresholds of dexamethasone destabilize CAR T-cell treatment efficacy

**DOI:** 10.1101/2021.10.01.462697

**Authors:** Alexander B. Brummer, Xin Yang, Eric Ma, Margarita Gutova, Christine E. Brown, Russell C. Rockne

## Abstract

Chimeric antigen receptor (CAR) T-cell therapy is potentially an effective targeted immunotherapy for glioblastoma, yet there is presently little known about the efficacy of CAR T-cell treatment when combined with the widely used anti-inflammatory and immunosuppressant glucocorticoid, dexamethasone. Here we present a mathematical model-based analysis of three patient-derived glioblastoma cell lines treated *in vitro* with CAR T-cells and dexamethasone. Advanced *in vitro* experimental cell killing assay technologies allow for highly resolved temporal dynamics of tumor cells treated with CAR T-cells and dexamethasone, making this a valuable model system for studying the rich dynamics of nonlinear biological processes with translational applications. We model the system as a nonautonomous, two-species predator-prey interaction of tumor cells and CAR T-cells, with explicit time-dependence in the clearance rate of dexamethasone. Using time as a bifurcation parameter, we show that (1) dexamethasone destabilizes coexistence equilibria between CAR T-cells and tumor cells in a dose-dependent manner and (2) as dexamethasone is cleared from the system, a stable coexistence equilibrium returns in the form of a Hopf bifurcation. With the model fit to experimental data, we demonstrate that high concentrations of dexamethasone antagonizes CAR T-cell efficacy by exhausting, or reducing the activity of CAR T-cells, and by promoting tumor cell growth. Finally, we identify a critical threshold in the ratio of CAR T-cell death to CAR T-cell proliferation rates that predicts eventual treatment success or failure that may be used to guide the dose and timing of CAR T-cell therapy in the presence of dexamethasone in patients.

**Author summary:** Bioengineering and gene-editing technologies have paved the way for advance immunotherapies that can target patient-specific tumor cells. One of these therapies, chimeric antigen receptor (CAR) T-cell therapy has recently shown promise in treating glioblastoma, an aggressive brain cancer often with poor patient prognosis. Dexamethasone is a commonly prescribed anti-inflammatory medication due to the health complications of tumor associated swelling in the brain. However, the immunosuppressant effects of dexamethasone on the immunotherapeutic CAR T-cells are not well understood. To address this issue, we use mathematical modeling to study *in vitro* dynamics of dexamethasone and CAR T-cells in three patient-derived glioblastoma cell lines. We find that in each cell line studied there is a threshold of tolerable dexamethasone concentration. Below this threshold, CAR T-cells are successful at eliminating the cancer cells, while above this threshold, dexamethasone critically inhibits CAR T-cell efficacy. Our modeling suggests that in the presence of high dexamethasone reduced CAR T-cell efficacy, or increased exhaustion, can occur and result in CAR T-cell treatment failure.

## Introduction

Chimeric antigen receptor (CAR) T-cell therapy is a rapidly advancing immunotherapy for the treatment of cancer. CAR T-cell therapy has demonstrated remarkable clinical outcomes in haematologic cancers, and this success has motivated efforts to advance CAR T-cell therapy for the treatment of solid tissue tumours, including the highly aggressive brain cancer glioblastoma (GBM) [1–4]. The prognosis for GBM following standard of care treatment of surgical resection, radiotherapy, and chemotherapy remains unacceptably low with most patients surviving less than 18 months [5]. CAR T-cell therapy may offer unrealized opportunities to improve outcomes for GBM based on the ability to engineer, expand, and adoptively transfer large numbers of tumor reactive T-cells. Our group and others are clinically evaluating CAR T-cells for the treatment of GBM, in which the therapeutic T-cells are delivered locoregionally [4, 6]. Our lead clinical program targets IL13R*α*2, a tumor associated antigen expressed by the majority of high-grade gliomas, including GBM [7, 8]. In early phase clinical trials, IL13R*α*2-CAR T-cells have shown encouraging evidence for antitumor bioactvitiy in a subset of patients [1, 9].

To further develop CAR T-cell therapy for the clinical treatment of GBM, it is essential to understand how CAR T-cells interact with commonly administered medications which may impact CAR T-cell efficacy. The anti-inflammatory synthetic glucocorticoid dexamethasone (Dex) is a ubiquitous medication for patients with GBM due to the propensity for brain tissue inflammation that accompanies tumor development in GBM, and the severity of the associated medical complications that accompanies inflammation. Dex is also commonly used to manage neurologic immune-related adverse events (irAEs) associated with CAR T-cells and other immunotherapies [10]. To study the effects of Dex on CAR T-cell proliferation, killing, and exhaustion, we extend mathematical models developed by us and others to study highly resolved temporal *in vitro* dynamics of patient-derived GBM cell lines under various concentrations of dexamethasone and CAR T-cells [11].

Recent work has demonstrated contradictory outcomes in the use of Dex for treating GBM. Specifically, the anti- and pro-proliferative effects of Dex on GBM have been shown to depend on cell type [12]. Furthermore, a previous proof-of-concept experiment demonstrated the ability of high Dex doses (5 mg/kg) to compromise successful CAR T-cell therapy in mice with xenograft GBM tumors, whereas lower doses (0.2-1 mg/kg) had limited effect on *in vivo* antitumor potency [8]. This data suggests a threshold at which Dex negatively impacts CAR T-cell therapy and reinforces the importance of mathematical modeling to infer and understand how Dex influences CAR T-cell therapy efficacy for GBM.

Mathematical modeling of CAR T-cells has demonstrated value in quantitatively characterizing tumor-immune cell dynamics. Compartmental models have been leveraged to enhance understanding of the underlying cancer biology. In such practices, variation in model complexity can be utilized to investigate either the myriad roles of T cell and tumor cell types [13–15] or the mathematical nature of the cell-cell interactions themselves [16, 17]. These same approaches can be naturally extended to inform and predict both pre-clinical and clinical applications of immunotherapies [18–21]. Most recently, such efforts have been applied to model and predict CAR T-cell therapies for leukemia [22, 23], glioblastoma organoids and solid brains tumors [11], and combination therapies with radiotherapy [24] and chemotherapy [25].

Previous work by us investigated CAR T-cell therapy for the treatment of glioblastoma (GBM) solid brain tumors. This work validated the principle components necessary for accurate predictions, specifically identifying: rates of GBM proliferation and cell killing, and CAR T-cell proliferation, exhaustion, and death. These factors were combined into a predator-prey system called CARRGO: Chimeric Antigen Receptor T-cell treatment Response in GliOma [11].

Here we extend this work by incorporating the Dex concentration as a new model parameter, and assume that it follows exponentially depleting pharmacokinetics. We posit that Dex has directly measurable effects on GBM proliferation and CAR T-cell death, and indirectly measurable effects on all other model parameters. We use our extended model to investigate the consequences of combination CAR T-cell and Dex therapy on three *in vitro* GBM cell lines. We establish an experimental protocol that measures treatment effects on GBM cell populations while co-varying initial CAR T-cell populations and Dex concentrations.

## Materials and methods

### Cell lines

Primary brain tumor (PBT) cell lines derived from GBM tumor resection tissue were derived as described in [1, 26]. All three cell lines come from male donors ages 43, 52, and 59 years old. As this study was focused on the interaction between Dex and CAR T-cells, cell lines were either selected based on the endogenous expression of IL13R*α*2 (PBT030 and PBT128) or engineered to express high levels of IL13R*α*2 (greater than 70%) by lentiviral transduction (PBT138) as described in [1, 11, 26]. Expression levels of IL13R*α*2 for each PBT cell line as determined by flow cytometry are shown in S1 Supporting Information. For IL13R*α*2-targeted CAR T-cell lines, healthy donor CD62L+ naive and memory T-cells were lentivirally transduced to express a second-generation of IL13R*α*2-targeting CAR as described in [8]. Summary information regarding cell lines can be found in Table 1.

**Table 1.** Experimental conditions.

| Tumor cell line<br>(% IL13Rα2) | CAR T-cells | Initial number of<br>tumor cells | Effector to Tar-<br>get (E:T) ratio | Dex concentrations<br>(μg/ml) |
| --- | --- | --- | --- | --- |
| PBT030 (97.97%) | IL13Rα2 BBζ | 10K-20K | 1:4, 1:8, 1:20 | 0, 10<sup>-4</sup>, 10<sup>-3</sup>, 10<sup>-2</sup>, 10<sup>-1</sup>, 1 |
| PBT128 (89.11%) | IL13Rα2 BBζ | 10K-20K | 1:4, 1:8, 1:20 | 0, 10<sup>-4</sup>, 10<sup>-3</sup>, 10<sup>-2</sup>, 10<sup>-1</sup>, 1 |
| PBT138 (99.53%) | IL13Rα2 BBζ | 10K-20K | 1:4, 1:8, 1:20 | 0, 10<sup>-4</sup>, 10<sup>-3</sup>, 10<sup>-2</sup>, 10<sup>-1</sup>, 1 |

Patient-derived brain tumor (PBT) lines with corresponding expression levels of IL13R $\alpha$ 2, CAR T-cell lines, initial number of cells, effector to target ratios, and Dex concentrations used in *in vitro* experiments.

### Experimental conditions

Cancer cell growth and treatment response was monitored with the xCELLigence cell analyzer system [27]. By correlating changes in electrical impedance with number of tumor cells adhered to electrode plates, a measurement of the cell population is reported every 15 minutes. Cell populations of both the tumor cells and CAR T-cells are reported in the non-dimensional units of Cell Index (CI), where 1 CI ≈ 10K cells. Previous work has demonstrated that cell index and cell number are strongly correlated [28, 29], including in the presence of CAR T-cell treatment [11]. Flow cytometry was used at the experiment endpoint to examine the validity of the cell index-cell number correlation in the presence of Dex, as well as count the non-adherent CAR T-cells. Tumor cells were seeded at 10K-20K cells per well and left either untreated, treated with only Dex, treated with only CAR T-cells, or treated with both Dex and CAR T-cells. All control and treatment conditions were conducted in duplicate, with treatments occurring 24 hours after seeding and followed for 6-8 days (144-192 hrs). CAR T-cell treatments were performed with E:T ratios of 1:4, 1:8, and 1:20. Dex treatment concentrations used were 10^−4^, 10^−3^, 10^−2^, 10^−1^, and 1 *μ*g/ml. The experiment design is diagrammed in Fig 1(**a**), and treatment conditions are presented in Table 1. A follow-up experiment was conducted to examine the potential for Dex-induced changes to tumor cell morphology using the IncuCyte live cell imaging system (Fig 2 and S1 Supporting Information). In this second experiment, E:T ratios of 1:20, 1:40, and 1:80 were used as the CAR T-cells had been engineered to be more efficacious. All other experimental conditions were held constant. See https://github.com/alexbbrummer/CARRGODEX for all experimental data.

**Fig 1.**
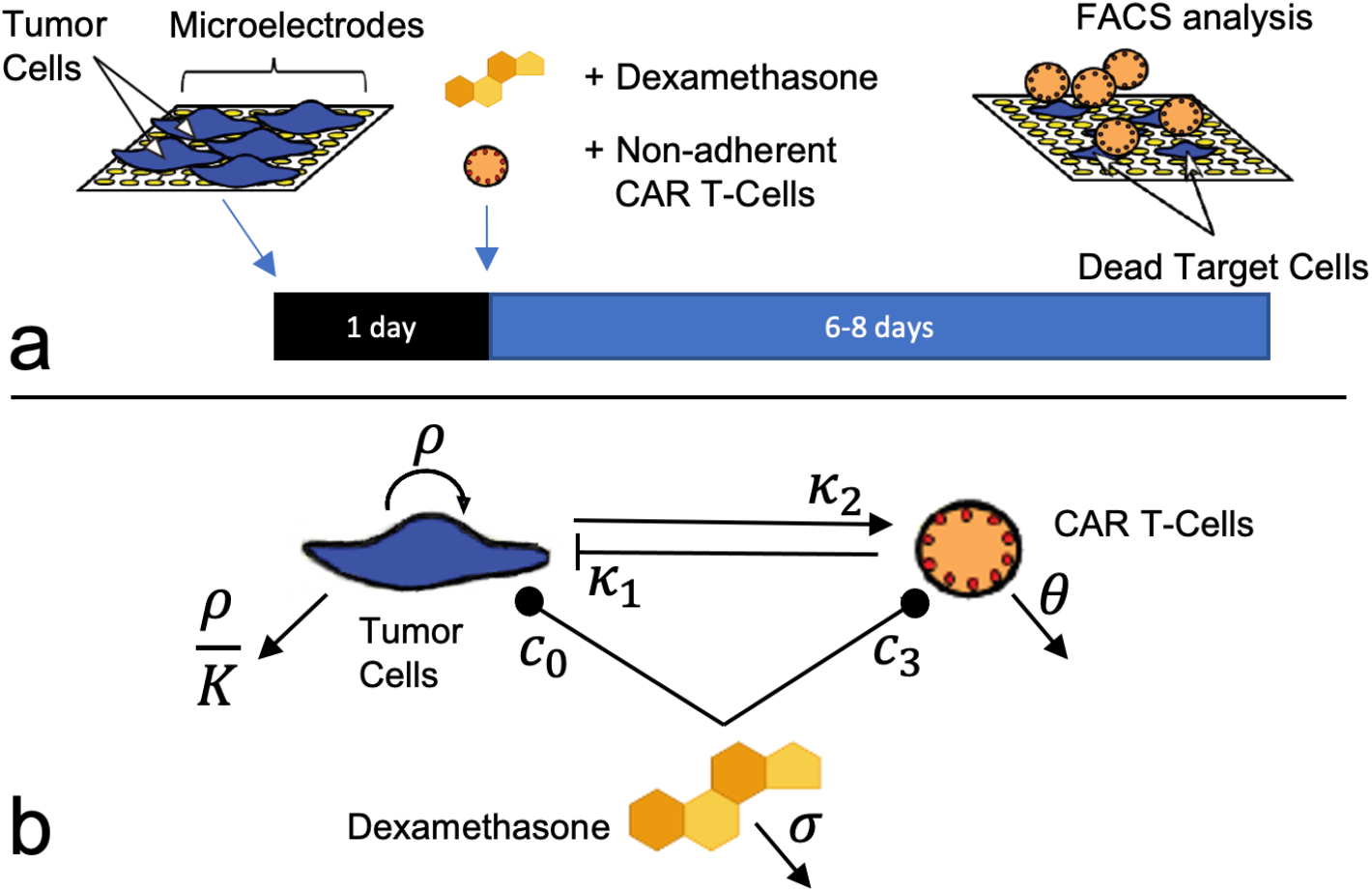
Diagram of *in vitro* experiments and mathematical model. (**a**) Experiments were conducted using a 96-well plate xCELLigence cell killing assay, where tumor cell adherence modulates electrical impedance. Dexamethasone and CAR T-cells were added simultaneously 24 hours following tumor cell plating, with observation proceeding for 6-8 days (144-192 hrs). CAR T-cells were counted at the experiment endpoint with flow cytometry analysis (FACS). (**b**) A mathematical model similar to a predator-prey system is used to model tumor cell growth, death, and interactions between tumor cells, CAR T-cells, and Dex. The compartmental model is translated into the system of equations in Eqs. (1)–(3).

**Fig 2.**
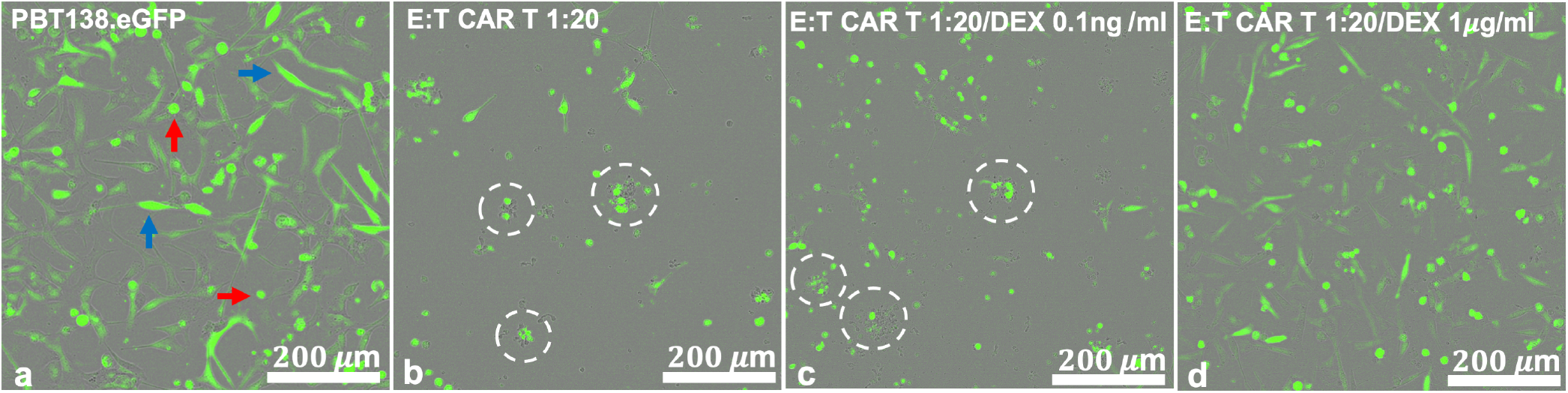
Incucyte live cell imaging of PBT138.eGFP cells 100 hours following treatment shows high Dex-induced reduction in CAR T-cell efficacy. (a) No Dex or CAR T-cell treatment (PBT138 culture). Visible are tumor cells of different morphology round (red arrows) and elongated (blue arrow), representing heterogeneous patient-derived glioma cell culture. (b) CAR T-cell only treatment with effector:target cell (E:T) ratio of 1:20. Visible are tumor apoptotic bodies that represent CAR T-cell killing (white circles). (c) Combined CAR T-cell (E:T = 1:20) and Dex (0.1 ng/ml) treatment. Tumor apoptotic bodies still visible (white circles) representing killing at low Dex. (d) Combined CAR T-cell (E:T = 1:20) and high Dex (1 *μ*g/ml) treatment showing lack of tumor aggregates and some increase in tumor cell numbers. White scale bars represent 200 *μ*m.

### Mathematical model

To model the interactions between the tumor cells, the CAR T-cells, and dexamethasone, we extend the predator-prey inspired CARRGO model from Sahoo et al. [11]. We use the principle of mass-action to model the effect of Dex on tumor and CAR T-cell populations, without an explicit assumption of a positive or negative effect of Dex on those cell populations. A compartmental representation of the model is presented in Fig 1(**b**), and all model variables and parameters are presented in Table 2. The tumor cell and CAR T-cell populations are modeled here in units of cell index (CI), a strongly correlated indicator of cell number that is produced by the xCELLigence cell killing assay measurement system [11, 28, 29]. Expressing the compartmental model as a system of equations,

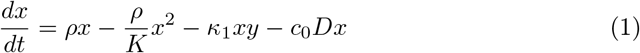

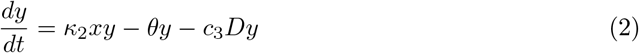

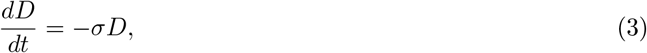

where *x* is the tumor cell population, *y* is the CAR T-cell population, and *D* is the concentration of dexamethasone. Although cell populations are often modelled in terms of cell number, here we use cell index (CI) to link model parameters with the experimental xCELLigence platform readout data. As determined in previous studies, cell index and cell number are strongly correlated, with a cell index of one equal to approximately 10,000 cells.

**Table 2.**
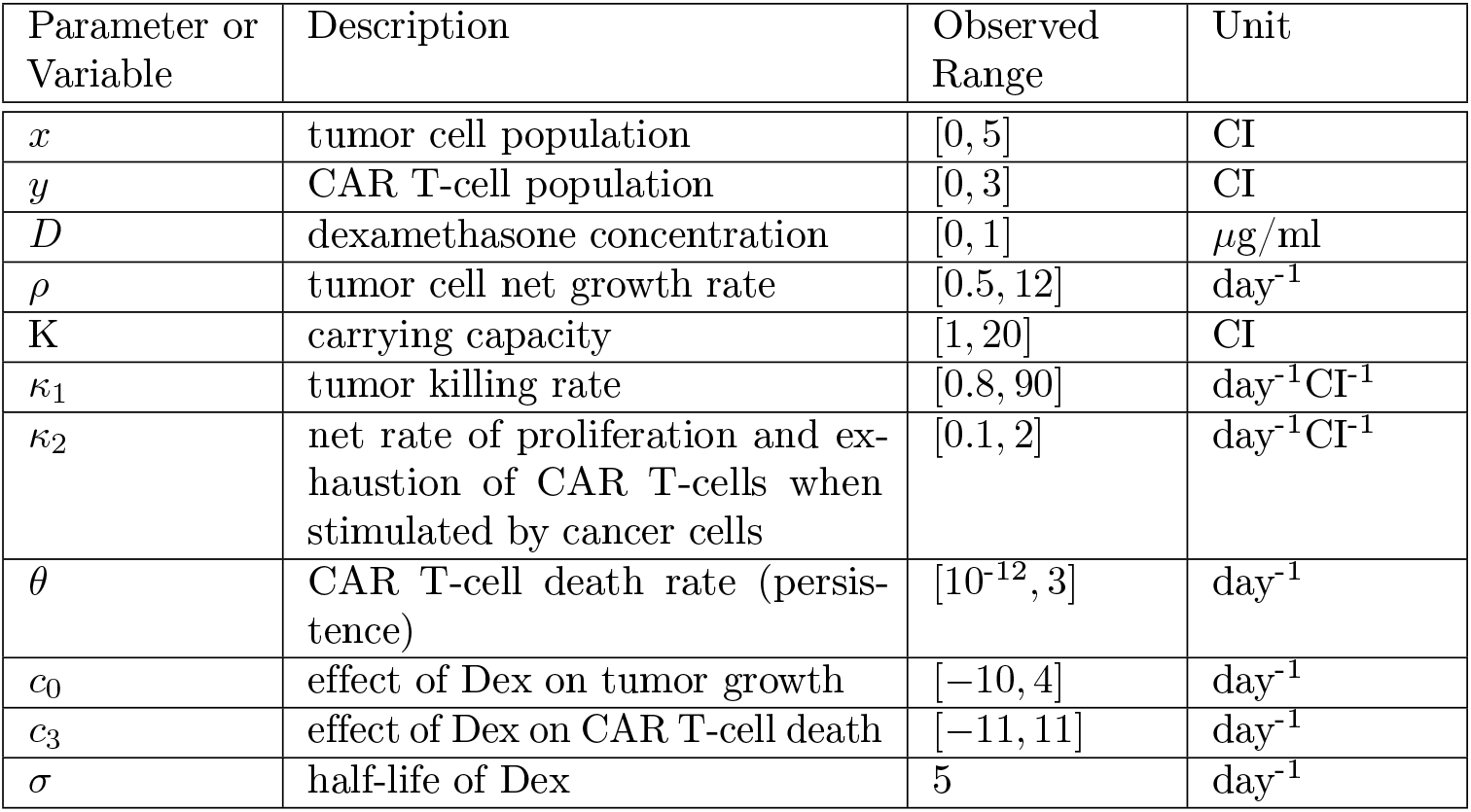
Mathematical model parameters and variables.

Pharmacokinetic studies report the plasma half-life of Dex as being approximately 200 minutes, resulting in 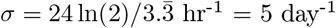 [30, 31]. We do not explicitly model the mechanism by which Dex is cleared from the system, which can be through cell uptake, evaporation, or absorption into the culture media. Here we simply assume the elimination of Dex is equivalent to the Dex plasma half-life. While the Dex interaction terms are explicitly subtracted from the population growth rates, we make no presumptions on the signs of the interaction constants, *c*_0_ and *c*_3_. This has the effect of allowing for both scenarios where Dex can be either anti-proliferative (i.e. *c*_0_, *c*_3_ < 0) or pro-proliferative (i.e. *c*_0_, *c*_3_ > 0) to either the CAR T-cells or tumor growth [12, 32].

We next convert this three-species, autonomous population model into a two-species, nonautonomous model. We formulate the model this way to study how the concentration of Dex influences the dynamical behavior and long-term stability of the CAR T-cell and tumor cell populations, which essentially considers time as a bifurcation parameter. The value of this approach is its utility in analyzing the stability of the tumor cell-CAR T-cell dynamics as time evolves, a perspective that bears more clinical relevance and simplicity than the exponentially decaying concentration of Dex.

Previous studies have utilized nonautonomous models to account for time-varying environmental conditions in generic predator-prey systems [33, 34], for pulsed patient preconditioning in combination CAR T-cell and chemotherapy [25], and in the analysis of pharmacokinetic-pharmacodynamic tumor growth models with time-dependent perturbations due to anticancer agents (see chapter 7 in [35]).

In the present model, we highlight the fact that the decaying dexamethasone concentration is modeled with a bounded and continuously differentiable function on the interval [0, ∞). This can be seen by separately solving Eq. (3) as *D*(*t*) = *D*_0_*e^−σt^*. Thus, stable solutions to the nonautonomous model will still converge to those of the autonomous model [35]. Upon substitution for *D*(*t*), we arrive at the following system of equations

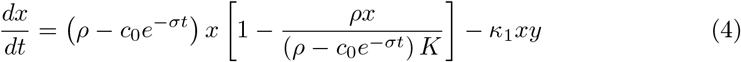

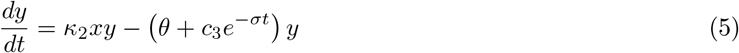

where we factored terms to reflect the anti/pro-proliferative potential of Dex, and re-scaled the constants *c*_0_ and *c*_3_. Letting *ρ*(*t*) = *ρ* − *c*_0_*e^−σt^*, *K*(*t*) = *ρ*(*t*)*K*/*ρ*, and *θ*(*t*) = *θ* + *c*_3_*e^−σt^*, our model takes the simplified form of

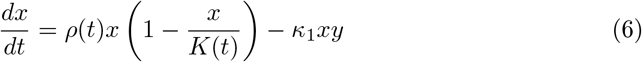

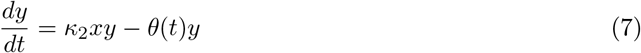

which is reminiscent of the original CARRGO model [11]. The definitions of *ρ*(*t*), *K*(*t*), and *θ*(*t*) demonstrate how the signs of the constants *c*_0_ and *c*_3_ can determine the anti/pro-proliferative effect of Dex on the tumor cells and CAR T-cells. Specifically, if *c*_0_ > 0, then Dex is anti-proliferative to the tumor cells, lowering the effective growth rate *ρ*(*t*) and carrying capacity *K*(*t*). If *c*_0_ < 0, then Dex is pro-proliferative to the tumor cells, raising the effective growth rate *ρ*(*t*) and carrying capacity *K*(*t*). On the other hand, if *c*_3_ > 0, then Dex is anti-proliferative to the CAR T-cells, increasing the effective death rate *θ*(*t*). If *c*_3_ < 0, then Dex is pro-proliferative to the CAR T-cells, decreasing the effective death rate *θ*(*t*).

### Parameter estimation

The fitting procedure used to estimate model parameters consisted of a combination of particle swarm optimization (PSO) and the Levenberg-Marquardt algorithm (LMA). PSO is a stochastic global optimization procedure inspired by biological swarming [36]. PSO has been used recently for parameter estimation in a variety of initial value problems across cancer research and systems biology [37–40]. These optimization procedures were used to minimize the weighted sum-of-squares error between measured and predicted tumor cell and CAR T-cell populations. PSO was used first to determine rough estimates of model parameters. This was followed by use of LMA to fine-tune parameter values.

Although the mathematical model has 8 parameters, we show that the model in Eqs. (6)–(7) is structurally identifiable from the experimental data. This allows for explicit measurement and inference of all the parameters in the model, including *c*_0_ and *c*_3_, and thus the effects of Dex on the tumor cells and CAR T-cells independently [41]. In particular, two replicates (wells) of tumor cells were grown untreated with CAR T-cells but with and without Dex treatments to independently identify the tumor growth rate, *ρ*, carrying capacity *K*, and the effect of Dex on the tumor growth rate and carrying capacity *c*_0_. Additionally, two replicates of tumor cells treated with CAR T-cells were conducted with and without Dex. This allowed us to independently identify the tumor killing rate with and without Dex, *κ*_1_, the CAR T-cell proliferation/exhaustion with and without Dex, *κ*_2_, the CAR T-cell death rate *θ*, and the effect of Dex on the CAR T-cell death rate *c*_3_. See Supplemental Information for identifiability analysis, and https://github.com/alexbbrummer/CARRGODEX for all experimental data and computational code to reproduce the model fitting, parameter estimates, and figures. All model fitting was performed using the programming language Python. All model fits were performed on averages of the experimental duplicates.

### Stability analysis

Prior work has demonstrated that conventional methods of stability analysis can be extended to nonautonomous models [35, 42]. As the concentration of dexamethasone is an exponentially decaying function, *D*(*t*) = *D*_0_*e^−σt^*, we can analyze the stability of Eqs. (4)–(5) as we would normally in an autonomous scenario. Despite this, in S1 Supporting Information we present a stability analysis of the 3×3 autonomous system for the coexistence equilibrium in the limit that the dexamethasone concentration decays to zero. This demonstrates that the eigenvalues of the two systems are effectively the same, with the only difference due to whether one expresses the eigenvalues in terms of the Dex concentration, D, or the precise form of its exponential decay, *D*_0_*e^−σt^*. Furthermore, we emphasize that the two systems converge on a time scale of the order of the decay constant, *σ*. Mathematically speaking, the Dex concentration never reaches zero, but on a more practical and physiological level, Dex clears after approximately 3-5 half lives, as presented later in the results.

With the simplified form of our CARRGO with Dex model in Eqs. (6)–(7), the equilibrium solutions are identified as *P*_1_ = (0, 0), *P*_2_ = (*K*(*t*), 0), and 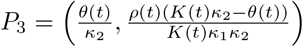 where (*x, y*) = (tumor cells, CAR T-cells). These solutions are referred to as ‘Death’, ‘Tumor Proliferation’, and ‘Coexistence’ respectively. Given the structure of the dynamical system in Eqs. (6)–(7), eigenvalue analysis shows that the ‘Death’ and ‘Tumor Proliferation’ equilibria are never stable solutions (see S1 Supporting Information). Interestingly, this does not preclude our ability to predict tumor death or proliferation. On the contrary, observed and measured tumor death and proliferation occur within the parameter space that defines the coexistence equilibrium. Careful examination of the coexistence equilibrium stability can elucidate this point.

In the ‘Coexistence’ scenario, the equilibrium is 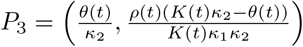. Importantly, the model parameters that determine the final tumor cell population are the ratio of the CAR T-cell death rate and the CAR T-cell proliferation/exhaustion after the Dex has cleared, *θ*/*κ*_2_. Thus, if either CAR T-cell death is low with respect to CAR T-cell proliferation, or CAR T-cell proliferation is high with respect to death, then *θ*/*κ*_2_ ≈ 0, and tumor death can occur as the coexistence equilibrium. We next examine how the conditions for stability depend on the model parameters, in particular the Dex concentration.

The eigenvalues of the Jacobian for the coexistence equilibrium are

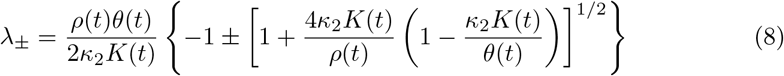

Recalling that *ρ*(*t*) = *ρ* − *c*_0_*e^−σt^* and 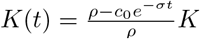, then 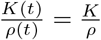. Substitution of the expressions for the time-dependent growth rate, carrying capacity, and death rate results in

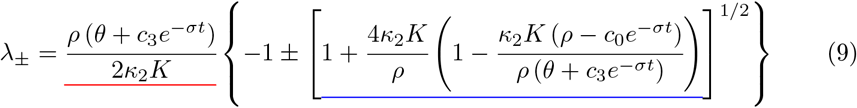

In Eq. (9), the term underlined in blue determines oscillatory behavior, while the term underlined in red determines the stability of the oscillatory states (spiraling in, spiraling out, or as a fixed limit cycle). By convention, only the parameters characterising the effects of Dex on tumor growth and CAR T-cell death, *c*_0_ and *c*_3_, can take on negative values. Thus, after the Dex has cleared, any oscillatory coexistence state will be stable. This consequence highlights the value of our decision to model the system as nonautonomous. In the event that Dex is pro-proliferative to the CAR T-cells such that *c*_3_ < −*θ*, then there will always be at least one positive eigenvalue (with or without oscillations), and the equilibrium will be temporarily unstable until the Dex has sufficiently cleared the system.

To determine the condition for oscillatory states, we require non-zero imaginary components of the eigenvalues, 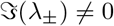, which results in the following condition

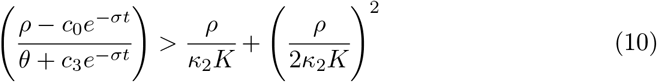

In Eq. (10) we can see that the existence of oscillations about the coexistence equilibrium are again determined by the relative sizes of *θ*(*t*) and *κ*_2_.

## Results

Experimental results of this study demonstrate that high concentrations of dexamethasone can attenuate tumor eradication even when CAR T-cell therapy would otherwise have been successful (Fig 2). This phenomenon was observed directly in cell line PBT128 (Fig 3), PBT138 (S1 Fig) and it can be inferred for cell line PBT030 between effector-to-target ratios 1:4 and 1:8 (S2 Fig and S3 Fig, respectively). A reduction in CAR T-cell efficacy is observed regardless of treatment success or failure at different effector-to-target ratios (E:T) for all three cell lines (Fig 4 and S1 Fig and S3 Fig). Importantly, our predator-prey model that incorporates Dex can predict these changes in CAR T-cell efficacy and connect them to key features of CAR T-cell function (e.g. proliferation, exhaustion, and death).

**Fig 3.**
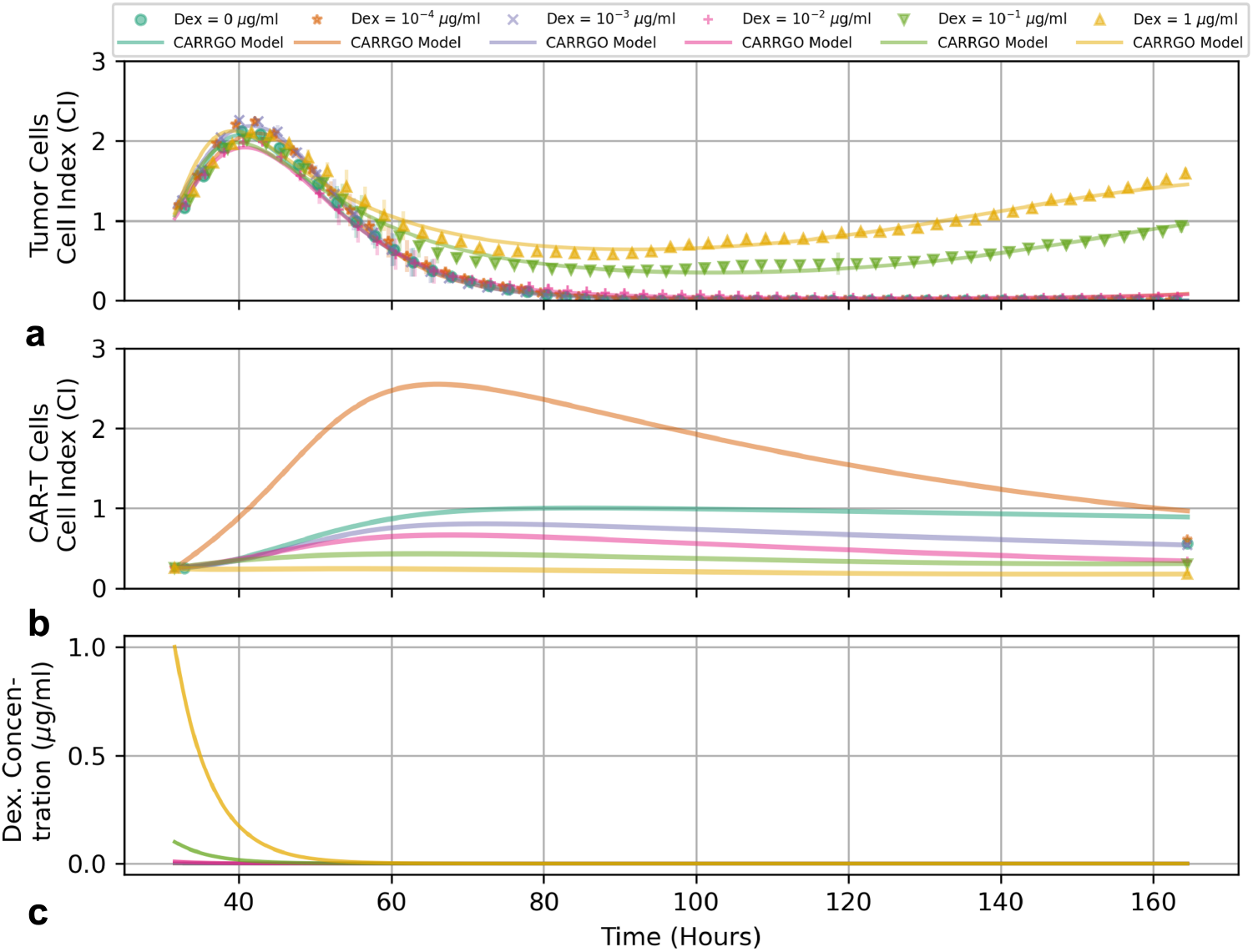
Graphs of measured and predicted (**a**) tumor cells, (**b**) CAR T-cells, and (**c**) Dex concentration over time for tumor cell line PBT128 with an initial effector-to-target ratio of 1:4. Temporal measurements of (**a**) tumor cells measured by xCELLigence cell index (CI) values and (**b**) CAR T-cell levels with initial and final measurements represented by symbols, and CARRGO model predictions are represented by lines. Experimental measurements for the tumor cell population are down-sampled by 1/10 for clarity. Colors and symbol types represent different initial Dex concentrations (see top legend). The progression of the tumor cell curves as initial Dex concentration increases demonstrate the effect of Dex to reduce CAR T-cell efficacy.

**Fig 4.**
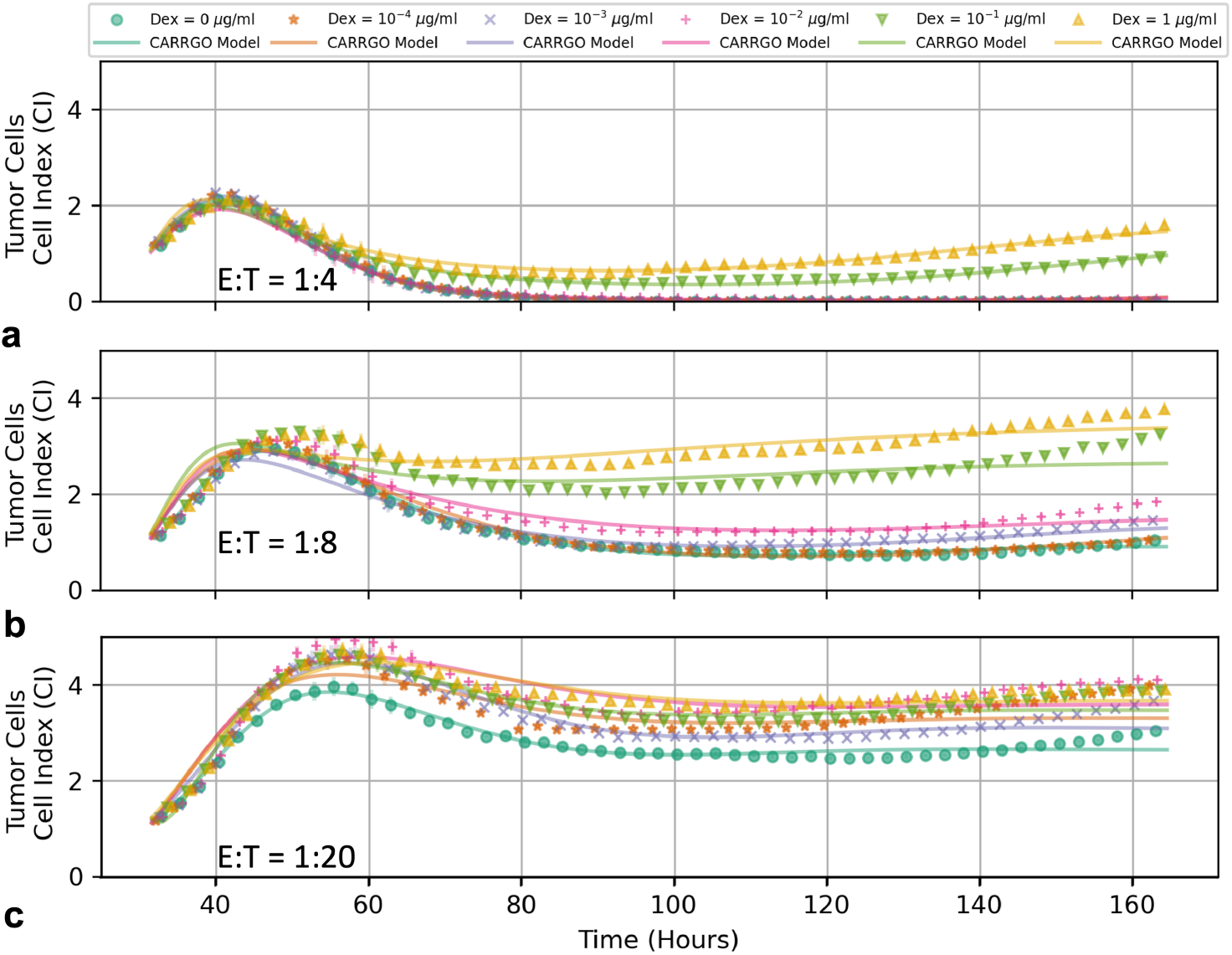
Time series of tumor cell populations for PBT128 across various CAR T-cell E:T ratios of 1:4 (**a**), 1:8 (**b**), and 1:20 (**c**). Colors and symbol types vary to reflect initial Dex concentrations (see top legend). Tumor persistence is seen to increase due to increasing initial Dex concentration regardless of the different starting CAR T-cell E:T ratios. Symbols represent measured data, while lines represent CARRGO model predictions. Experimental measurements for the tumor cell population are down-sampled by 1/10 for clarity.

### Dexamethasone antagonizes CAR T-cell efficacy

Our experimental results suggest that Dex acts to antagonize CAR T-cell treatment efficacy in a dose-dependent manner, resulting in persistent tumor cell growth. In Fig 3 we see that final populations of tumor cells for line PBT128 increase as a function of increasing initial Dex concentration. Specifically, the system dynamics range from complete tumor cell death for initial Dex concentrations of 0.0 *μ*g/ml, 1 × 10^−4^ *μ*g/ml, 1 × 10^−3^ *μ*g/ml, and 1 × 10^−2^ *μ*g/ml to tumor progression for initial Dex concentrations of 1 × 10^−1^ *μ*g/ml, and 1 *μ*g/ml. We also see a noted decrease in initial CAR T-cell growth as the initial Dex concentration is increased.

The loss of CAR T-cell treatment efficacy as a result of Dex can be observed across all CAR T-cell E:T ratios. In Fig 4 we see how increasing the initial Dex concentration continually increases the final tumor population when compared to the Dex control for the PBT128 cell line. Interestingly, the extent to which pseudo-regression occurs for the medium and low dose CAR T-cell groups is noticeably diminished as Dex increases.

Also notable are the cases of medium initial CAR T-cells (E:T = 1:8) and high initial Dex (0.1 *μ*g/ml −1 *μ*g/ml), where the final tumor cell population has surpassed the initial pseudo-progression peak. Flow cytometry measurements of these high Dex concentrations in the absence of CAR T-cell treatment for PBT128 demonstrate that the xCELLigence Cell Index metric begins to overestimate tumor cell number (S2 Fig). This effect was also observed in PBT030 (again in the absence of CAR T-cells), but not PBT138, and led to our omission of the Dex-only treatments in this analysis.

### Dexamethasone induced destablization of coexistence

Analysis of the coexistence eigenvalue stability helps to elucidate the effect that Dex has on the system dynamics. Fig 5 presents bifurcation diagrams for two different experimental scenarios with the same initial CAR T-cell population but different initial Dex concentrations. Fig 5**a** shows the coexistence eigenvalues as functions of time for the experimental conditions of high initial CAR T-cells (E:T = 1:4) and a low initial Dex concentration of 1 × 10^−3^ *μ*g/ml in which productive tumor cell death occurred. Fig 5**b** shows the coexistence eigenvalues as functions of time for the experimental conditions of high initial CAR T-cells (E:T = 1:4) and a high initial Dex concentration of 1 × 10^−1^ *μ*g/ml and corresponds to tumor cell progression. To illustrate how the time-dependence of the eigenvalues influences the system dynamics, streamplots of Eqs. (6)–(7) are presented as insets for each experimental scenario, with experimentally measured tumor cell index values and model-inferred CAR T-cell index vales represented by the black dots.

**Fig 5.**
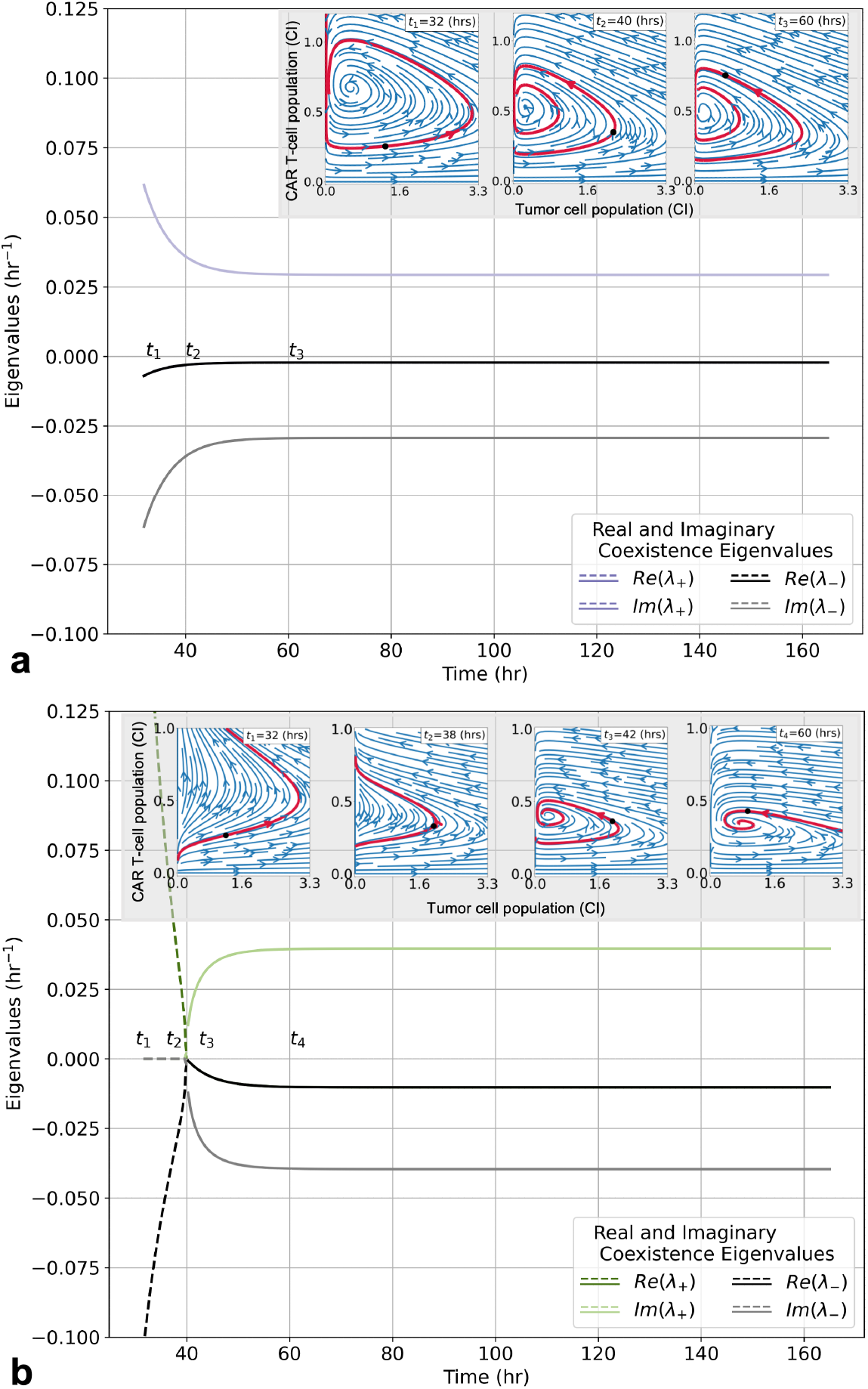
Stability analysis of coexistence equilibria for CARRGO model with Dex under the experimental conditions of tumor cell line PBT128 with an initial CAR T-cell E:T ratio of 1:4. Initial Dex concentrations are 1 × 10^−3^ *μ*g/ml (**a**) and 1 × 10^−1^ *μ*g/ml (**b**). Bifurcation diagrams (main graphs) demonstrate how Dex modulates the temporal dynamics of the coexistence equilibria. Inset graphs show phase space diagrams demonstrating transient state of coexistence equilibria as time evolves and Dex clears. In (**a**) the initial Dex concentration of 1 × 10^−3^ *μ*g/ml is too small to facilitate exhaustion of the CAR T-cells, thus the coexistence equilibrium is a stable spiral. However, the equilibrium position is still seen to translate through the phase space, and the predicted trajectory deform, while Dex is clearing. In (**b**) the initial Dex concentration of 1 × 10^−1^ *μ*g/ml is sufficiently large enough to facilitate exhaustion of the CAR T-cells, thus the coexistence equilibrium is an unstable fixed point (dashed lines) until a sufficient level of Dex has cleared and the system returns to a stable spiral (solid lines). Unlike the lower initial Dex concentration scenario in (**a**), the coexistence equilibrium has translated, and the predicted trajectory has narrowed, such that the tumor population no longer reaches a value of zero, resulting in tumor progression.

In the scenario with an initial Dex concentration of 1 × 10^−3^ *μ*g/ml (Fig 5 **a**), the coexistence equilibrium begins as a stable spiral for the duration of the Dex clearance and the remainder of the experiment. As the Dex clears, the real and imaginary components of the eigenvalues decrease in magnitude. These temporal changes in the eigenvalues shift the location and shape of the system trajectory, as shown in the figure inset. In particular, throughout the times *t*_1_ = 32 hrs and *t*_2_ = 40 hrs, as the Dex is still clearing, the system is predicted and observed to oscillate about the changing coexistence equilibrium *P*_3_. Initially, and throughout times *t*_1_ and *t*_2_, the real component of the eigenvalue is large enough to facilitate in-spiraling. By *t*_3_ = 60 hrs, effectively all of the Dex has cleared, and the phase space trajectory is soon to pass through a zero in tumor cell population, terminating the dynamics.

In the scenario with a higher initial Dex concentration of 1 × 10^−1^ *μ*g/ml (Fig 5 **b**), the coexistence equilibrium begins as an unstable fixed point (represented by the dashed lines during times *t*_1_ = 32 hrs and *t*_2_ = 38 hrs). After twice the half-life of Dex (≈7 hrs), a Hopf bifurcation occurs and the system transitions through a limit cycle (observable at time *t* ≈40 hrs) and into a stable spiral (represented by the solid lines during time *t*_3_ = 42 hrs). When the system is in an unstable state, the instantaneous trajectory predicted by Eqs. (6)–(7) show pseudo-progressive growth. As in the previous scenario with 1 × 10^−3^ *μ*g/ml of Dex, the system enters a stable spiral by the time all of the Dex has cleared (*t*_4_ = 60 hrs). However, in this scenario, once the Dex has fully cleared the system the predicted trajectory no longer passes through a zero in the tumor cell population.

### Tumor cell killing and CAR T-cell exhaustion

To examine the effect of Dex on total tumor cell killing we compare tumor cell growth trajectories for CAR T-cell only treatment, and combined CAR T-cell and Dex treatments in Fig 6. For the combined treatment scenarios, we focus on experimental conditions at the threshold of treatment success and treatment failure, and along a Dex-gradient of treatment failure (Fig 6 **a**, **c**, and **e** for cell lines PBT030, PBT128, and PBT138, respectively). Accompanying each growth trajectory are barplots of the inferred model parameters (Fig 6 **b**, **d**, and **f**) which help to identify mechanistically how Dex interacts separately with the tumor cells and CAR T-cells. We chose not to examine Dex only treatments as flow cytometry measurements indicated a loss in the strength of the correlation between xCELLigence cell index and flow cytometry-measured cell number in these treatment scenarios [28, 29]. The correlation was observed to be maintained in combined treatment scenarios, with correlation coefficients of = 0.97 in cell line PBT138, 0.70 in cell line PBT128, and 0.30 in cell line PBT030 (S1 Supporting Information).

**Fig 6.**
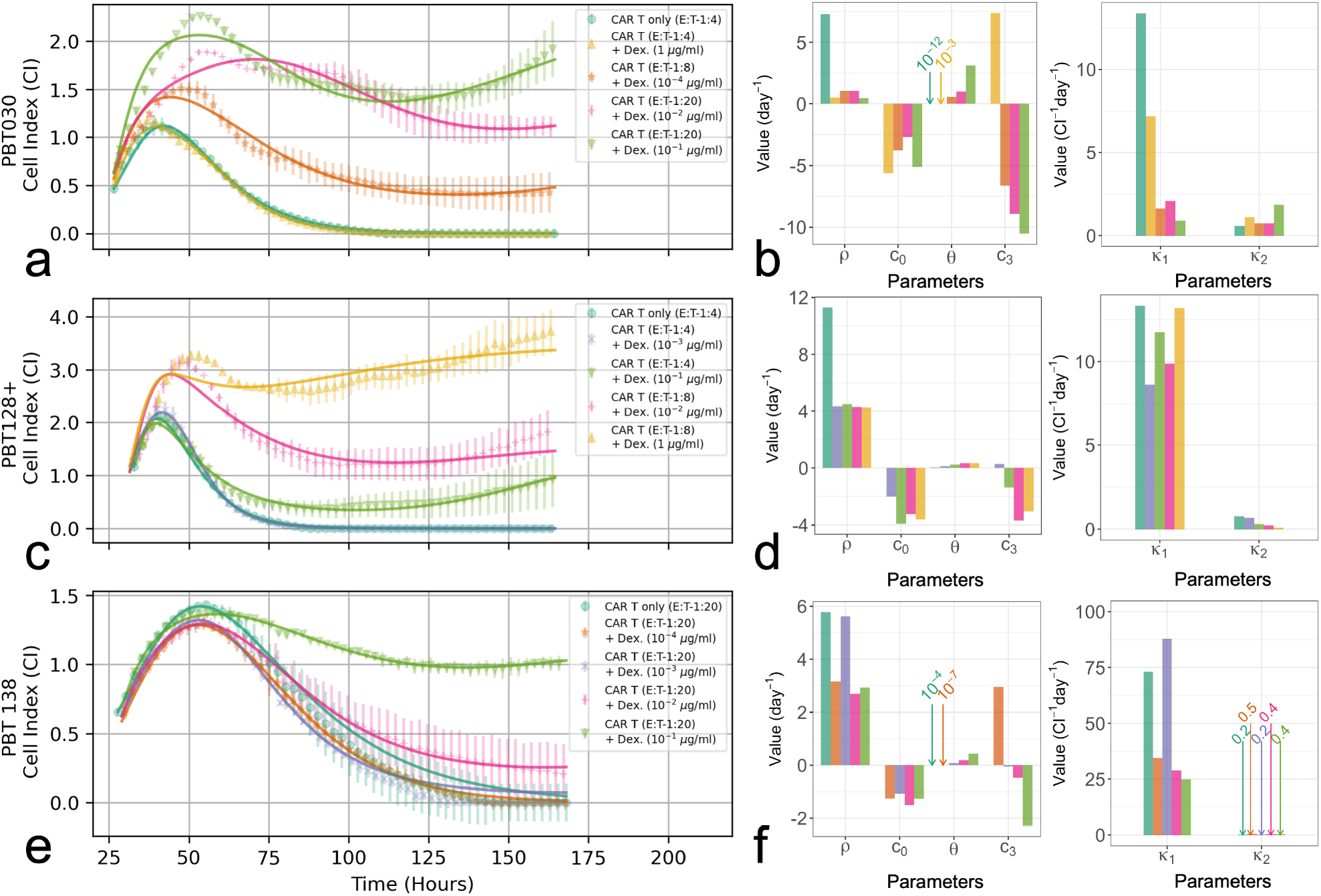
Comparison of tumor cell measurements and CARRGO model fits (**a**, **c**, and **e**), and CARRGO model parameters (**b**, **d**, and **f**) for CAR T-cell only treatments, and combination CAR T-cell and Dex treatments for each tumor cell line studied. For the combined treatment scenarios, we focus on experimental conditions at the threshold of treatment success and treatment failure, and along a Dex-gradient of increasing tumor progression. For PBT030 (**a**, **b**) the conditions at the threshold of treatment success are E:T = 1:4 and 1 *μ*g/ml of Dex, and E:T = 1:8 and 10^−4^ *μ*g/ml for treatment failure. For PBT128 (**c**, **d**) the conditions at the threshold of treatment success are E:T = 1:4 and 10^−3^ *μ*g/ml of Dex, and E:T = 1:4 and 10^−1^ *μ*g/ml for treatment failure. For PBT138 (**e**, **f**) the conditions at the threshold of treatment success are E:T = 1:20 and 10^−4^ *μ*g/ml of Dex, and E:T = 1:20 and 10^−3^ *μ*g/ml for treatment failure. Symbols represent average of two replicates, and error bars represent sample ranges. Note that experimental measurements presented are downsampled by 1/10 for clarity.

From the perspective of our mathematical model, the effect of Dex on the CAR T-cells is to reduce efficacy by inducing exhaustion. In going from treatment success to failure, the CAR T-cell death rate, *θ*, increases for all cell lines. Furthermore, for cell lines PBT030 and PBT138 the tumor cell killing, *κ*_1_, decreases while CAR T-cell proliferation, *κ*_2_, remains fixed, while in cell line PBT128 the tumor cell killing, *κ*_1_, remains fixed while CAR T-cell proliferation, *κ*_2_, decreases. These shifts dramatically increase the predicted coexistence equilibrium for the tumor cell population, given as *θ*/*κ*_2_. Next is the effect of Dex on the CAR T-cell death rate, *c*_3_, which switches from positive to negative between success and failure across all cell lines. This shift suggests that during treatment success there is high turnover of CAR T-cells due to the Dex-induced increase in CAR T-cell death and the cancer cell stimulated CAR T-cell proliferation. After Dex clears, CAR T-cell death returns to a small rate resulting in treatment success. In the failure scenario Dex again promotes CAR T-cell growth, *c*_3_ < 0, yet increases CAR T-cell exhaustion by reducing the size of *κ*_2_. Our interpretation of these combined effects is that Dex results in CAR T-cell exhaustion.

### Predicting treatment success or failure

Analysis of treatment success and failure across all experimental conditions identifies an essential threshold, *T*_0_, for the predicted tumor cell equilibrium population. As shown in Fig 7, we observe an approximate threshold of *T*_0_ ≈ 0.4 CI such that for values of *θ*/*κ*_2_ < *T*_0_, total tumor cell death occurs after a brief period of pseudo-progression. Alternatively, for values of *θ*/*κ*_2_ > *T*_0_, tumor cell persistence occurs after pseudo-progression and pseudo-response. Importantly, for the cell lines PBT128 and PBT138, we can see a transition from treatment success to treatment failure at fixed levels of initial CAR T-cells (0.25CI for PBT128 and 0.05CI PBT138) and as initial Dex concentration increases from 10^−2^ to 10^−1^ *μ*g/ml. For tumor cell line PBT030, the transition from treatment success to failure occurs primarily as a result of changes in the initial number of CAR T-cells administered, with E:T ratios of 1:4 resulting in success, and 1:8 resulting in failure.

**Fig 7.**
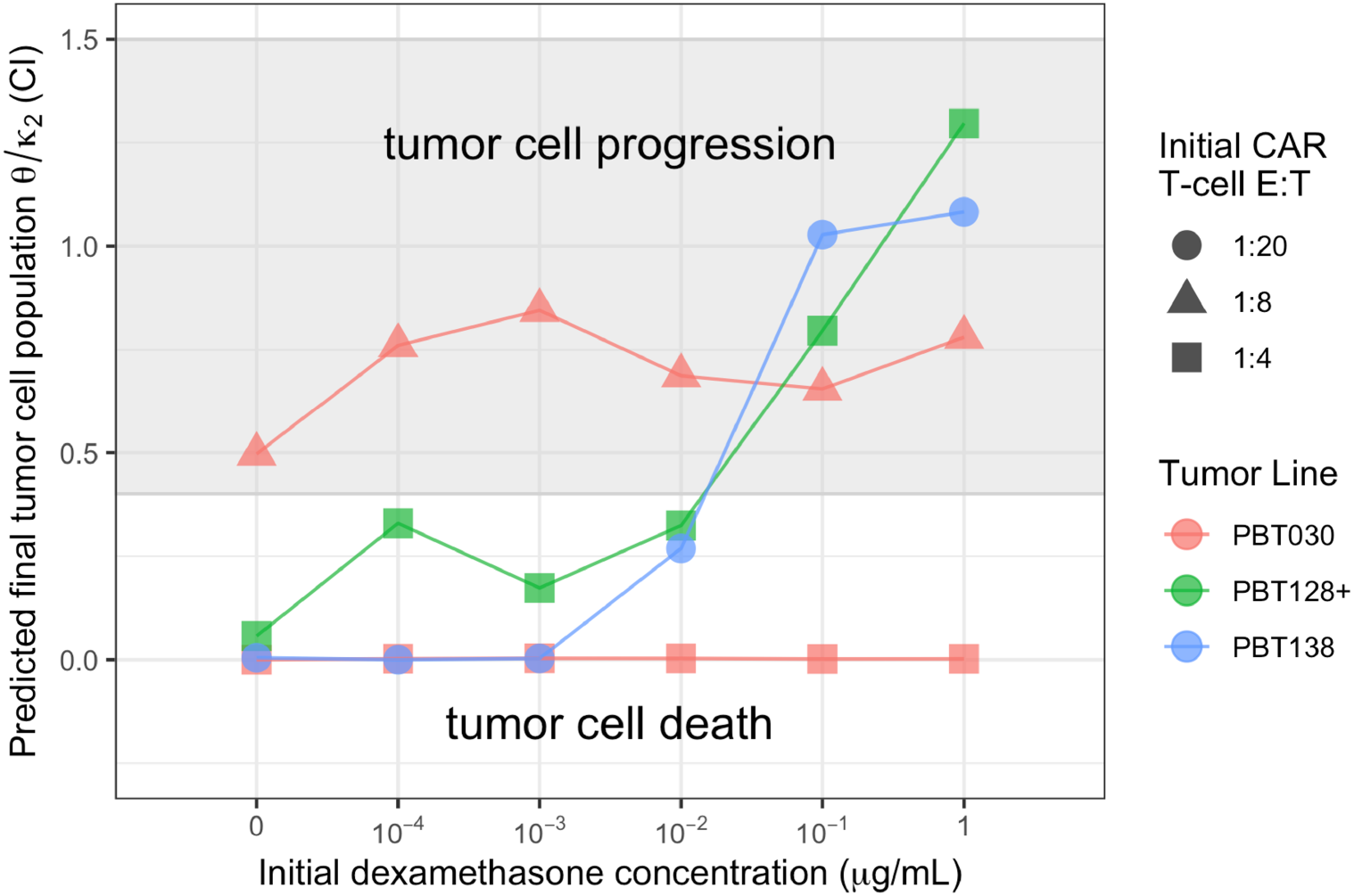
The ratio of CAR T-cell death (*θ*) to CAR T-cell proliferation/exhaustion (*κ*_2_) rates predict CAR T-cell treatment success (tumor cell death) or failure (tumor cell progression). We observed that a ratio of *θ*/*κ*_2_ ≈ 0.4 CI as the predicted final tumor cell population serves as a threshold for observed tumor progression or death. The threshold of *θ*/*κ*_2_ ≈ 0.4 CI was consistent across all three tumor cell lines (denoted by color), CAR T-cell E:T ratios (denoted by shape), and initial Dex concentrations (denoted by location along the horizontal axis). Conditions not shown are PBT030 with an E:T ratio of 1:20 which all resulted in tumor cell progression, PBT128 with an E:T ratio of 1:8 and 1:20 which all resulted in tumor cell progression, and PBT138 with E:T ratios of 1:8 and 1:4 which all resulted in tumor cell death. See S4 Fig for all E:T ratios.

## Discussion

In this work we demonstrate how mathematical modeling can be leveraged to identify and quantify how the commonly used anti-inflammatory synthetic glucocorticoid dexamethasone may undermine CAR T-cell treatment efficacy in glioblastoma.

Our modeling identifies that Dex treatment destabilizes the coexistence equilibrium and forces the system into a new equilibrium state upon Dex clearance (Fig 5). We predict that this process is a result of a Dex-induced increase in CAR T-cell proliferation, *c*_3_ < −*θ*, followed by an increase in CAR T-cell death, *θ* increasing, and either a decrease or fixation of cancer cell stimulated proliferation of CAR T-cells, *κ*_2_ decreasing or approx. constant. We interpret these combined effects as facilitating CAR T-cell exhaustion (Fig 6). These responses are manifest in a cycle of pseudo-progression, pseudo-regression, and a final stage of tumor progression. Importantly, we identify a threshold on the ratio of CAR T-cell death to CAR T-cell proliferation/exhaustion rates, (*θ*/*κ*_2_ ≈ 0.4 CI), that appears to predict successful tumor eradication (*θ*/*κ*_2_ < 0.4 CI), or proliferation (*θ*/*κ*_2_ > 0.4 CI). We find that this threshold is valid across all three PBT cell lines, initial CAR T-cell populations, and Dex concentrations (Fig 7).

In modeling this system, we chose to use a nonautonomous (explicit in time) approach in order to assess the dynamical stability of the system as a function of Dex concentration. By treating time as a bifurcation parameter, variations in system stability and predicted phase space trajectories can be visualized (Fig 5). This approach facilitates understanding of treatment success and failure due to an overabundance of Dex and in the context of stability analysis. Specifically, in scenarios where treatment succeeded, as in Fig 5(**a**), the coexistence equilibrium remained stable throughout the duration of Dex clearance. On the other hand, in scenarios where high levels of Dex led to treatment failure and tumor outgrowth, as in Fig 5(**b**), the coexistence equilibrium was initially unstable until sufficient Dex cleared from the system.

Insight gained from studying tumor-CAR T-cell dynamics can aid in understanding how Dex levels compromise CAR T-cell efficacy. Destabilization of the coexistence equilibrium is driven by changes to the experimentally derived model parameters *θ*, *κ*_2_, and *c*_3_ (Fig 6 and Eq. (10), which represent the death rate of CAR T-cells, the proliferation or exhaustion of the CAR T-cells, and the effect of Dex on the CAR T-cell death rate, respectively. These changes are interpreted as Dex promoting tumor cell growth and CAR T-cell growth at early times (due to *c*_3_ < −*θ*), yet once the Dex has cleared the CAR T-cells become exhausted, no longer proliferating enough to keep up with natural death or facilitate tumor cell killing. CAR T-cell exhaustion, indicated by a decrease in *κ*_2_, is a primary cause of tumor progression as determined by the increase in the predicted final tumor cell population, *θ*/*κ*_2_ (Fig 6). Importantly, this Dex-induced shift from CAR T-cell proliferation to exhaustion is highlighted by the differences in phase space trajectories between treatment success and treatment failure presented in Fig 5.

A notable feature of the mathematical model is the fact that it captures a wide range of dynamics observed in multiple experimental conditions [11]. This is despite its relative simplicity compared to other mathematical models of immunotherapies [13, 17, 43, 44]. A recent commentary regarding predator-prey like models, including the model presented here, is the possibility of oscillating solutions which are unlikely to be observed in patients [45, 46]. We note that the coexistence equilibrium is accompanied with phase-space trajectories that accurately describe experimental data. This includes scenarios of treatment success and tumor death (*x* = 0), allowing for informative and quantitative biological inference. Furthermore, we highlight that each model parameter can be uniquely identified from our measured data, as supported by our structural identifiability analysis (S1 Supporting Information) [41]. This allows for the deconvolution of dynamics and parameters not otherwise accessible from the cell killing assay.

Several possible extensions of our model exist and are worth consideration in future analyses. A common extension is to generalize the interaction between tumor cells and CAR T-cells to a higher order Holling Type form [13, 17, 43]. Generally, the Holling Type II and III interactions are used to model changes in cell-cell interactions, notably predator-prey handling time and density-dependent behavior. Additionally, where the CARRGO model combines CAR T-cell proliferation and exhaustion into one parameter, *κ*_2_, other approaches may incorporate a second T-cell type altogether [20, 45], a population of macrophages [44], or explicitly accounting for the pharmacodynamic and pharmacokinetics of CAR T-cell dynamics [47]. Interestingly, recent theoretical work has shown that a two T-cell type predator-prey model with Holling Type I interactions can, in the appropriate limits and conditions, reduce to a single T-cell type predator-prey model with a Holling Type II interaction [43]. While such model extensions can be enlightening, our approach aims to balance model complexity with the dimensionality and resolution of the experimental data.

### Limitations and simplifications

Several limitations and simplifications were made in the course of this work that naturally suggest follow up studies. Although these studies were informative to assess the direct effect of Dex on CAR T-cell effector function, there is potential to construct models with many interacting immune populations. The fact that the xCELLigence cell killing assay is an *in vitro* system lacking an immune system naturally limits the model complexity that is experimentally accessible. Related to this is the clearance rate of the dexamethasone, assumed here to have a fixed value of approximately 200 minutes. In patient populations, some level of variation in the clearance rate is to be expected due to physiological differences [30, 31]. Future work examining how variation in the clearance rate affects treatment success would be of interest. Furthermore, the fact that T-cells are non-adherent to the cell killing assay precludes proposed models that require high temporal resolution of the T-cell dynamics. Presently, our experimental protocol includes only two datapoints for the CAR T-cells: the initial and final timepoints. Although this results in a boundary value problem from a mathematical point of view and uniquely determined solution for the CAR T-cell population, an immediate benefit from a modeling perspective would be high temporal resolution measurement of CAR T-cell dynamics similar to the tumor cells. Another simplification is that this modeling framework does not include spatial variations in tumor cell, CAR T-cell, or Dex density. Due to the highly structured nature of the brain and heterogeneity of glioblastoma tumors, including hypoxia, necrosis, and extensive invasion through the brain, spatial considerations may be important. Finally, a growing subject of importance is understanding sex and age-based differences in the immunological responses of patient derived cell lines, and how those difference translate to an individual level in clinical applications [48]. In this work, all GBM cell lines were derived from male patients of a similar age (43, 52, and 59 years old), suggesting that future *in vitro* work would benefit from including a greater diversity of patients across both age and sex.

### Potential applications and clinical relevance

Translating our findings to clinical applications requires refining understanding of the treatment success or failure threshold in terms of clinically accessible information for the treatment of GBM. In previous studies, we established that low doses of subcutaneous injections of Dex (0.2-1 mg/kg) had limited effect on *in vivo* antitumor potency in orthotopic murine models of GBM, whereas high doses of Dex (5 mg/kg) significantly compromise successful CAr T-cell therapy.

The goal of this study was to extend these findings by modeling a wide-range of *in vitro* Dex levels to better predict CAR T-cell responses in the presence of Dex. Our finding evaluating IL13R*α*2-CAR T-cells suggest that *in vitro* Dex concentrations between 10-100 ng/mL would correspond to this treatment failure threshold. While *in vivo* Dex concentrations locally in the brain and tumor microenvironment are difficult to measure and depend on the blood brain barrier, vasculature, and brain fluid flow, it has been reported that patients receiving oral administration of 7.5 mg of Dex result in serum Dex concentrations ranging from 2.5 to 98.1 ng/mL (median 61.6 ng/mL) within 1 to 3 hours [49], a range encompassing the treatment threshold defined in this study. In our phase 1 clinical trials evaluating CAR T-cells for GBM, Dex is limited to 6 mg per day in an effort to balance the clinical utility of Dex for reducing tumor-associated edema and immune related inflammation during CAR T-cell therapy, while at the same time maximizing CAR T-cell treatment efficacy, and in light of this data we are continuing to evaluate the clinical impact of Dex on therapeutic activity.

Another important consideration is the role of Dex on endogenous immune responses. In a study evaluating neoantigen vaccine therapy for GBM, the generation of polyfunctional neoantigen T-cell responses was severely compromised in patients receiving Dex during T-cell priming [50]. Importantly, the interplay between CAR T-cell therapy and endogenous immune responses has been shown to positively contribute to the treatment success of CAR T-cell therapy [51–53]. Thus, the impact of Dex on host immune responses, which was not evaluated in this study, will be an important future consideration when assessing the effect of Dex on CAR T-cell therapy.

Further, as advances in personalized medicine continue to develop patient-specific treatment plans, it is important to consider how many CAR T-cells are required for effective treatment in addition to how much Dex. This question is essential for designing patient specific adaptive therapies and in use of clinical decision support software. Furthermore, it requires knowledge of the spatial extent of individual tumors, the *in vivo* spatial heterogeneity of CAR T-cells and dexamethasone concentrations, and patient response and tolerance to timed-drug delivery.

One can also consider varying the concentration and timing of the high dexamethasone dosages to still provide therapeutic levels of Dex yet avoid compromising CAR T-cell efficacy. Previous theoretical work analysing pulsed drug delivery shows promise for this alternative approach [19]. Yet still, another treatment strategy could be patient preconditioning with dexamethasone followed with delayed, and perhaps pulsed, CAR T-cell delivery. Recent simulated studies investigating preconditioning with chemotherapy [25] or targeted radionuclide therapy [24] followed with CAR T-cells suggests that combination pretreatment and time-delay approaches have clinical value, in particular for providing therapeutic dosages at lower total concentrations. Based on the duration of observable changes to the phase-space trajectory in Fig 5, we suggest a time-delay of 2-3 dexamethasone half-lives.

While adaptive therapy protocols have yet to be fully implemented in CAR T-cell treatment plans, data driven methods such as clinical decision support systems and other machine learning inspired approaches have been proposed for patient monitoring [54]. Here, algorithms are trained on historical patient treatment data in an effort to assess the likelihood that new patients will develop cytokine release syndrome (CRS) or immune effector cell-associated neurotoxicity syndrome (ICANS) as a result of CAR T-cell therapy. Typical management plans involve, among other things, the use of corticosteroids such as dexamethasone. In this context, our results emphasize the need to better resolve the threshold of treatment success in combining CAR T-cells and Dex given that severe and unexpected complications can occur when trying to predict a treatment response that involves nonlinear drug interactions.

An essential component to understanding and predicting combination CAR T-cell and dexamethasone treatment success in clinical scenarios is the spatial extent and heterogeneity of tumors and the spatial variation of CAR T-cell and dexamethasone concentrations. While advances in medical imaging and patient-specific treatment planning are aiding this effort, equally important is the development of spatially-dependent models that can accurately account for observed variation. Recent work in this direction has shown promise, demonstrating the ability of mathematical models to combine genotypic evolution with spatial aggregation to describe heterogeneous tumor growth [55].

## Conclusion

In this work, we present an analysis of experimental data designed to untangle the interaction between glioblastoma cancer cells, CAR T-cells, and the anti-inflammatory glucocorticoid, dexamethasone. We examined three different human derived primary brain tumor glioblastoma cell lines and found that dexamethasone can act to exhaust CAR T-cells leading to tumor outgrowth, thereby undermining treatment success. In cases of extreme dosing, this results in complete treatment failure and tumor progression. Our use of a nonautonomous, explicitly time-dependent predatory-prey model to characterize the interactions demonstrates that dexamethasone acts to destabilize the coexistence equilibrium between CAR T-cells and tumor cells.

Furthermore, we observe that the predicted coexistence equilibrium population for the tumor cells, defined as the ratio of the CAR T-cell death rate to the CAR T-cell proliferation/exhaustion rate, serves as an experimental threshold for treatment success or failure. This work has important implications for future clinical applications of combination therapy using CAR T-cell and dexamethasone, as well as demonstrates the value of using nonautonomous models for pharmacodynamics.

## Supporting information

**S1 Supporting Information. Supplementary Text.** Contains analysis of regression between xCELLigence cell index and flow cytometry cell number, xCELLigence time series and IncuCyte imaging of dexamethasone only treatments, expression levels of IL13R*α*2 in primary brain tumor cell lines, stability analysis for the autonomous 3 × 3 coexistence equilibrium and for the nonautonomous 2 × 2 ‘Death’ and ‘Tumor Proliferation’ equilibria, structural identifiability of the CARRGO with Dex model, and methods for parameter estimation by particle swarm optimization.

**S1 Fig. Data and model fits for PBT138 with E:T=1:20.** Graphs of tumor cells, CAR T-cells, and Dex concentration over time for tumor cell line PBT138 with an initial effector to target ratio of 1:4. Temporal measurements of tumor cell population and the initial and final CAR T-cell measurements are represented by symbols, and CARRGO model predictions are represented by lines. Colors and symbol types vary to reflect initial Dex concentrations (see top legend). The progression of the tumor cell curves as initial Dex concentration increases demonstrate the effect of Dex to reduce CAR T-cell efficacy. In particular, CAR T-cell treatment is successful at low Dex initial Dex concentrations (0, 10^−4^, and 10^−3^ *μg/ml*) and fails at higher initial Dex concentrations (10^−2^, 10^−1^, and 1 *μg/ml*), resulting in tumor cell progression. Experimental measurements for the tumor cell population are down-sampled by 1/10 for clarity.

**S2 Fig. Data and model fits for PBT030 with E:T=1:4.** Similar graphical information as S1 Fig. presented for tumor cell line PBT030 with an initial effector-to-target ratio of 1:4. For all initial Dex concentrations, treatment success is observed, resulting in complete tumor death.

**S3 Fig. Data and model fits for PBT030 with E:T=1:8.** Similar graphical information as S1 Fig. presented for tumor cell line PBT030 with an initial effector-to-target ratio of 1:8. For all initial Dex concentrations, treatment failure is observed, resulting in tumor cell progression that generally increases with increasing initial Dex concentrations.

**S4 Fig. Treatment success/failure threshold.** The ratio of CAR T-cell death (θ) to CAR T-cell proliferation/exhaustion (*κ*_2_) rates predict CAR T-cell treatment success (tumor cell death) or failure (tumor cell progression). We observed that a ratio of *θ*/*κ*_2_ ≈ 0.4 CI as the predicted final tumor cell population serves as a threshold for observed tumor progression or death. The threshold of *θ*/*κ*_2_ ≈ 0.4 CI was consistent across all three tumor cell lines (denoted by color), CAR T-cell E:T ratios (denoted by shape), and initial Dex concentrations (denoted by location along the horizontal axis).

## Data Availability

All data and code used to perform the analyses and generate figures are available on a Github repository at https://github.com/alexbbrummer/CARRGODEX.

## Acknowledgments

This manuscript is dedicated to the memory of Xin (Cindy) Yang; she will always be remembered for her important scientific contributions, caring nature and smile.

Research reported in this publication was supported by the California Institute of Regenerative Medicine (CIRM) under CLIN2-10248 and the National Cancer Institute of the National Institutes of Health under grant numbers R01CA254271 (CB), R01NS115971 (CB, RR), and P30CA033572. The content is solely the responsibility of the authors and does not necessarily represent the official views of the National Institutes of Health. The authors would also like to acknowledge the support of the Marcus Foundation.

## Author Contributions

**Conceptualization:** CEB, RCR. **Data Curation:** ABB, XY, EM. **Formal Analysis:** ABB. **Funding Acquisition:** CEB, RCR. **Investigation:** ABB, XY, EM, MG, CEB, RCR. **Methodology:** ABB, XY, EM, MG, CEB, RCR. **Project Administration:** CEB, MG, RCR. **Resources:** CEB, MG, RCR. **Software:** ABB **Supervsion:** CEB, MG, RCR. **Validation:** ABB, XY, EM, MG, CEB, RCR. **Visualization:** ABB, RCR. **Writing—Original Draft Prepration** ABB. **Writing—Review & Editing:** ABB, EM, MG, CEB, RCR.

