## Supplementary material for "Dose-dependent thresholds of dexamethasone destabilize CAR T-cell treatment efficacy": S1 Supporting Information

### S1 Supporting Information for Dose-dependent thresholds of dexamethasone destabilize CAR T-cell treatment efficacy

#### 1 Correlation between xCelligence cell index and flow cytometry cell number

An essential feature of the experimental system is the ability of the xCelligence cell killing assay to accurately measure changes in cell number. The xCelligence system uses adherent cells to modify the electrical resistance, essentially converting cell number into an electric potential difference measured as voltage. The xCelligence cell index has been shown to correlate strongly with cell number—measured with conventional flow cytometry [1, 2]. Prior work examining CAR T-cell killing dynamics in the absence of Dexamethasone has demonstrated that the correlation between these metrics remains strong [3]. To support our ability to use the xCelligence cell index values to accurately characterize the system dynamics, we conducted flow cytometry measurements at the conclusion of our experiments to validate the correlation strength between cell index and cell number. These values are presented in Fig. SA, and the regression coefficients are presented in Table SB.

**Table SA. Regression coefficients for xCelligence cell index against flow cytometry cell count.**

| Regression data | PBT030 | PBT128 | PBT138 | All data |
|---|---|---|---|---|
| Includes E:T = 0 | 0.07 | 0.64 | 0.95 | 0.38 |
| Does not include E:T = 0 | 0.30 | 0.70 | 0.97 | 0.77 |

We found that cell index and cell number were highly correlated in cell line PBT138, and moderately correlated in cell line PBT128. Interestingly, PBT030 showed poor correlation between cell index and cell number. This was driven specifically by a group of outliers in which the treatment consisted only of Dexamethasone, or where the effector to target ratio was E:T = 0. Noting that these same treatment conditions led to

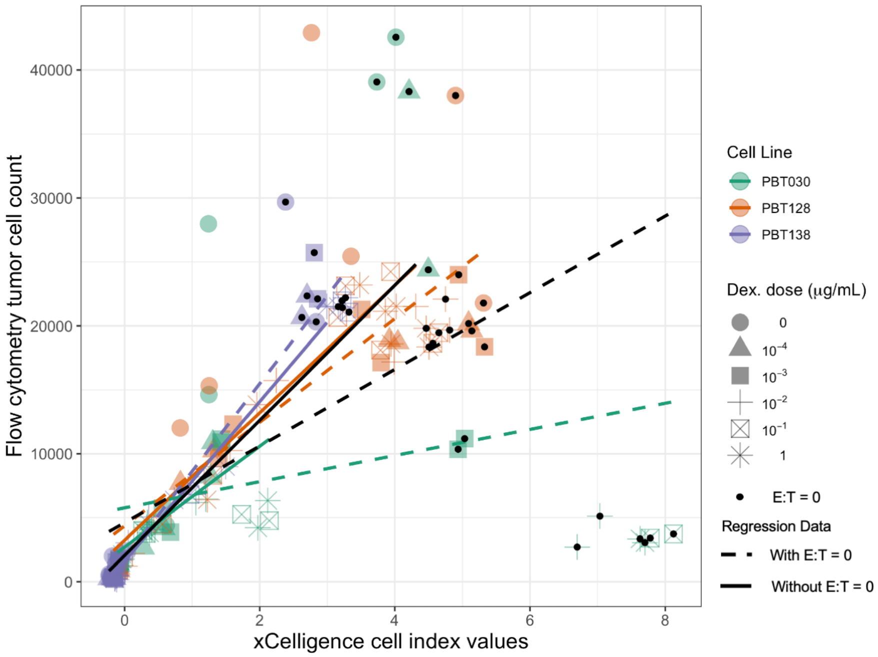


**Fig SA. Correlation between cell number and cell index.** Graphed are flow cytometry cell counts and the cell index values measured with the xCelligence system from the start and end points of the first experiment. Regressions are performed for individual cell lines and aggregated across all Dexamethasone concentrations. In black are regressions for all cell lines combined. Regression statistics are provided in Table SB.

outlier groupings in the other cell lines, we chose to omit all treatment scenarios in which only Dexamethasone was applied, or in which the effector to target ratio was E:T = 0. Fig. SA shows improved agreement between cell line-specific correlations as a direct result of this omission. Regression coefficients reported in Table SB further support this decision.

#### 2 Expression levels of IL13R$\alpha$2 in primary brain tumor cell lines

**Table SB. IL13R$\alpha$2 expression levels.**

| PBT Cell line | % IL13$\alpha$2 | MFI | ISO MFI | Diff MFI |
|---|---|---|---|---|
| PBT030 | 98.0 | 39173 | 254 | 38919 |
| PBT128 | 89.1 | 1804 | 240 | 1563 |
| PBT138 | 99.5 | 71433 | 6408 | 65024 |

Median flourescence intensity (MFI) of IL13R$alpha$2 expression on PBT lines when compared with staining using corresponding control -PE conjugated antibodies.

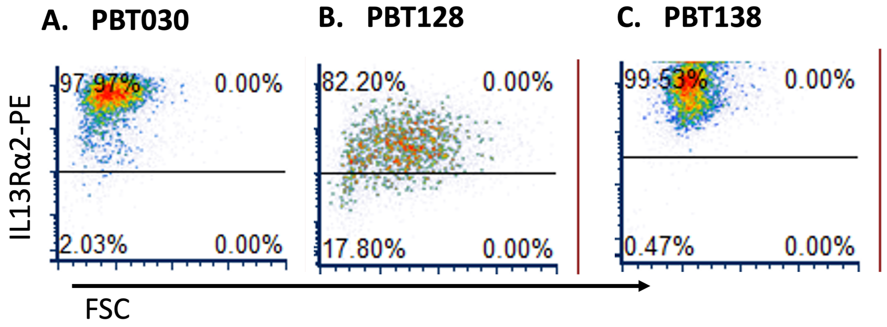


**Fig SB.** (A-C) flow cytometry dot plots demonstrating IL13R$\alpha$2 positive populations, labeled with IL13R$\alpha$2-PE conjugated antibodies. (D) Median florescence intensity (MFI) of IL13R$\alpha$2 expression on PBT lines when compared with staining using corresponding isotype control -PE conjugated anitbodies.

### 3 xCELLigence time series and IncuCyte imaging of dexamethasone only treatments

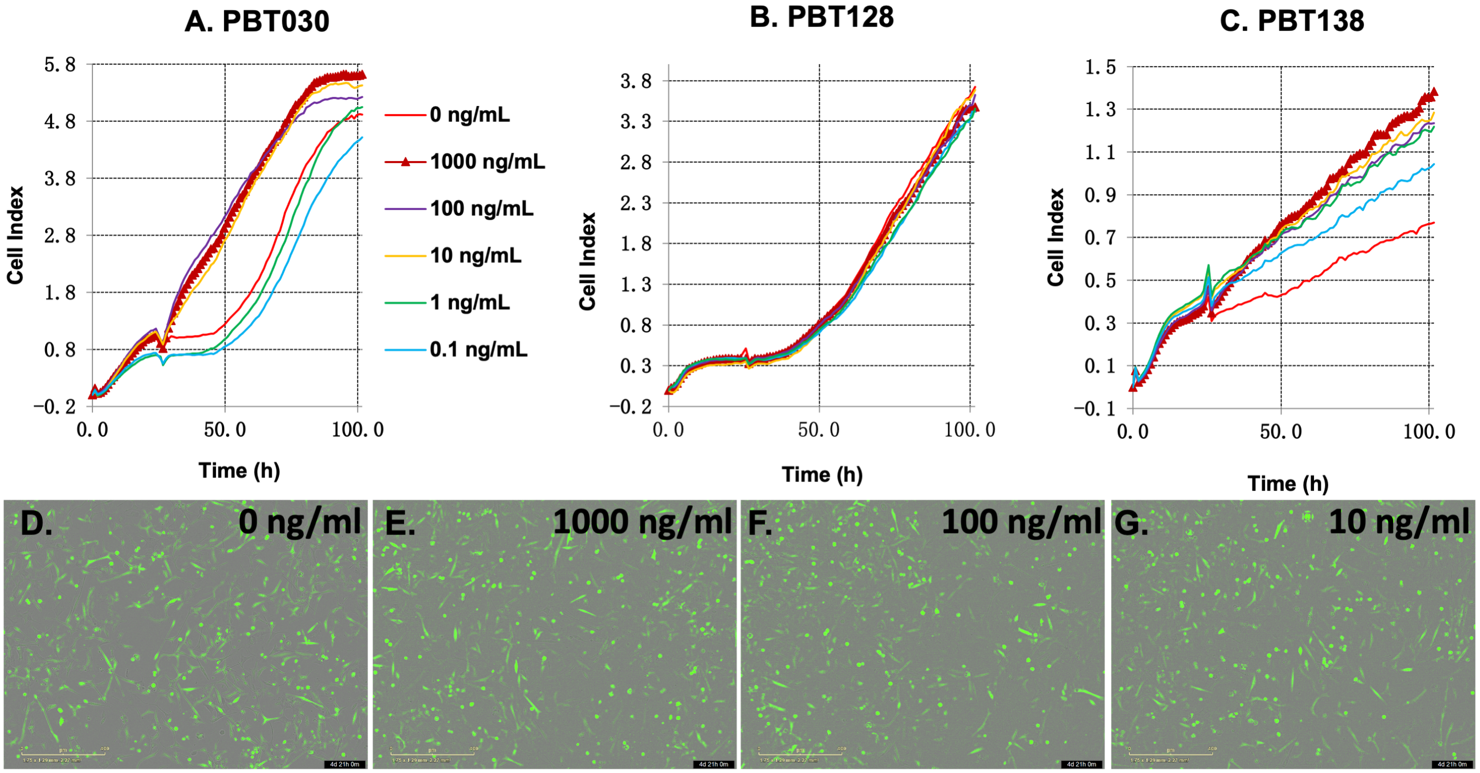


**Fig SC.** (A-C) xCELLigence measurements of PBT glioma lines treated with only concentrations of dexamethasone ranging from 0 to 1000 ng/ml (no effector CAR T-cells present). (C) PBT138 showing tumor proliferation dependent on Dex dosage. Data in (A-C) represents two independent experiments performed in duplicates. (D-G) IncuCyte images of PBT138 glioma line expressing eGFP, indicating no significant changes in morphology upon treatment with dexamethasone.

### 4 The coexistence equilibrium in the autonomous CARRGO model with Dex

We present here a linear stability analysis of the autonomous version of the CARRGO model with Dex. From the main text, this model is defined as,

$$\frac{dx}{dt} = \rho x - \frac{\rho}{K}x^2 - \kappa_1 xy - c_0 Dx \tag{S1}$$

$$\frac{dy}{dt} = \kappa_2 xy - \theta y - c_3 Dy \tag{S2}$$

$$\frac{dD}{dt} = -\sigma D, \tag{S3}$$

Following rearrangement to simplify future algebra, we have,

$$\frac{dx}{dt} = (\rho - c_0 D)x\left(1 - \frac{\rho x}{K(\rho - c_0 D)}\right) - \kappa_1 xy \tag{S4}$$

$$\frac{dy}{dt} = \kappa_2 xy - (\theta + c_3 D)y \tag{S5}$$

$$\frac{dD}{dt} = -\sigma D, \tag{S6}$$

Finally, we define the following variables, $\tilde{\rho} = \rho - c_0 D$, $\widetilde{K} = \tilde{\rho}K/\rho$, and $\tilde{\theta} = \theta + c_3 D$. Substituting these variables results in,

$$\frac{dx}{dt} = \tilde{\rho}x\left(1 - \frac{x}{\widetilde{K}}\right) - \kappa_1 xy \tag{S7}$$

$$\frac{dy}{dt} = \kappa_2 xy - \tilde{\theta} \tag{S8}$$

$$\frac{dD}{dt} = -\sigma D, \tag{S9}$$

As the Dex concentration is assumed to follow the continuously differentiable exponential decay, and we are interested in anayzing the dynamics of the system during the clearance of Dex, we consider the coexistence equilibrium in the limit that the Dex concentration approaches zero. This allows us to keep the Dex concentration variable as nonzero while performing stability analysis. Thus, the coexistence equilibrium for the autonomous system has coordinates $x^* = \tilde{\theta}/\kappa_2$, $y^* = \tilde{\rho}(\widetilde{K}\kappa_2 - \theta)/(\widetilde{K}\kappa_1\kappa_2)$, $D^* = D$. The Jacobian for this system is expressed as,

$$J = \begin{bmatrix} r\tilde{h}o - \frac{2\tilde{\rho}x^*}{\widetilde{K}} - \kappa_1 y^* & -\kappa_1 x^* & \frac{\partial \dot{x}}{\partial D}\Big|_* \\ \kappa_2 y^* & \kappa_2 x^* - \tilde{\theta} & \frac{\partial \dot{y}}{\partial D}\Big|_* \\ 0 & 0 & -\sigma \end{bmatrix} \tag{S10}$$

Upon evaluation of the coexistence equilibrium coordinates, the eigenvalues of the Jacobian for the coexistence equilibrium are,

$$\lambda_\sigma = -\sigma \tag{S11}$$

$$\lambda_\pm = \frac{\rho(\theta + c_3 D^*)}{2K\kappa_2} \left\{ -1 \pm \left[ 1 + \frac{4K\kappa_2}{\rho} \left( 1 - \frac{K\kappa_2(\rho - c_0 D^*)}{\theta + c_3 D^*} \right) \right]^{\frac{1}{2}} \right\} \tag{S12}$$

As $\sigma$ represents the half-life of the decaying Dex, it is always positive, and thus $\lambda_\sigma$ is always negative. Furthermore, the eigenvalues $\lambda_\pm$ take on the exact form as those from the nonautonomous coexistence equilibrium in the main text. However, in the autonomous model, variation in the equilibrium occurs due to the decaying Dex, $D$, where as in the nonautonomous model it is due to the evolution of time. This can be seen following substitution of $D = D_0 e^{-\sigma t}$. From this point on, details regarding how stability varies as a function of the model parameters as time evolves takes the same form as that presented in the main text for the nonautonomous model.

### 5 The 'death' and 'tumor proliferation' equilibria in the nonautonomous CARRGO model with Dex

In general, the Jacobian for the CARRGO model with Dex is expressed as

$$J(t) = \begin{bmatrix} \rho(t) - \frac{2\rho(t)}{K(t)} x - \kappa_1 y & -\kappa_1 x \\ \kappa_2 y & \kappa_2 x - \theta(t) \end{bmatrix} \tag{S13}$$

In the 'Death' scenario, the equilibrium is $P_1 = (0, 0)$ and the Jacobian becomes

$$J_1(t) = \begin{bmatrix} \rho(t) & 0 \\ 0 & -\theta(t) \end{bmatrix} \tag{S14}$$

with eigenvalues $\sigma_1 = \rho(t) = \rho - c_0 e^{-\sigma t}$ and $\sigma_2 = -\theta(t) = -\theta - c_3 e^{-\sigma t}$. As $\rho$ is always positive, then there is at least one positive eigenvalue, and the 'Death' scenario is an unstable equilibrium.

In the 'Tumor Proliferation' scenario, the equilibrium position $P_2 = (K(t), 0)$ and the Jacobian becomes

$$J_2(t) = \begin{bmatrix} -\rho(t) & -\kappa_1 K(t) \\ 0 & \kappa_2 K(t) - \theta(t) \end{bmatrix} \tag{S15}$$

with eigenvalues $\sigma_1 = -\rho(t) = -\rho + c_0 e^{-\sigma t}$ and
$\sigma_2 = \kappa_2 K(t) - \theta(t) = \left(\frac{\rho - c_0 e^{-\sigma t}}{\rho}\right) \kappa_2 K - \theta - c_3 e^{-\sigma t}$. As $\kappa_2$, $K$, and $\rho$ are always positive, there is always at least one positive eigenvalue, and the 'Tumor Proliferation' scenario is an unstable equilibrium.

### 6 Structural identifiability

An important aspect of the extended CARRGO model presented in Eqs. (5)-(6) is addressing the nature and extent of structural identifiability. Specifically, given the anticipated measurable quantities, are the model parameters uniquely identifiable or not? To answer this question, we assume the following:

- $\rho$, $K$, and $c_0$ are known. These parameters can be uniquely determined by fitting measurements for untreated and DEX-only treated tumor cell growth.
- The population of CAR T-cells at time $t = 0$ hrs is known, and is defined as $y_0$.
- The population of tumor cells at time $t = 0$ hrs is known, and is defined as $x_0$.
- The functions describing the tumor cell population, $x(t)$, and the CAR T-cell popluation are continuously differentiable.
- The following derivatives of the tumor cell population at time $t = 0$ hrs are known: $x_1 = dx/dt|_{t=0}$, $x_2 = d^2x/dt^2|_{t=0}$, $x_3 = d^3x/dt^3|_{t=0}$, $x_4 = d^4x/dt^4|_{t=0}$. This assumption comes from the fact that with appropriate fitting of the measured data, higher order derivatives can be calculated.

What remains is demonstrating that the remaining model parameters, $\kappa_1$, $\kappa_2$, $\theta$, and $c_3$, are identifiable. Starting with Eq. (5), we can solve for the tumor cell killing rate, $\kappa_1$, by setting time $t = 0$ hrs. This results in the following

$$\kappa_1 = \frac{(\rho - c_0)}{y_0} \left[1 - \frac{\rho x_0}{(\rho - c_0)K}\right] - \frac{x_1}{x_0 y_0} \tag{S16}$$

As all terms are assumed known in Eq. (S16), this demonstrates identifiability of $\kappa_1$. For the remaining parameters, we highlight that higher derivatives of the CAR T-cell population at time $t = 0$ hrs are needed. These values can be accessed by examining higher order derivatives of Eq. (5). For example, the second derivative with respect to time of the tumor cell population, evaluated at time $t = 0$ hrs, is given by,

$$x_2 = (\rho + c_0\sigma)x_0 + (\rho - c_0)x_1 - \frac{2\rho x_0 x_1}{K} - \kappa_1 x_1 y_0 - \kappa_1 x_0 y_1 \tag{S17}$$

Solving for the first derivative of the CAR T-cell population, $y_1$, results in,

$$y_1 = \frac{(\rho + c_0\sigma)}{\kappa_1} + \frac{(\rho - c_0)x_1}{\kappa_1 x_0} - \frac{2\rho x_1}{K\kappa_1} - \frac{x_1 y_0}{x_0} - \frac{x_2}{\kappa_1 x_0} \tag{S18}$$

Similar steps can be taken to identify the values of $y_2$, and $y_3$ using expressions for the higher derivatives of $x_3$, and $x_4$. From this point on, we assume that the values of $y_2$ and $y_3$ are known.

To identify the remaining model parameters, $\kappa_2$, $\theta$, and $c_3$, we now consider Eq. (6) and the following higher order derivatives,

$$y_1 = \kappa_2 x_0 y_0 - (\theta + c_3) y_0 \tag{S19}$$

$$y_2 = \kappa_2(x_1 y_0 + x_0 y_1) + c_3\sigma y_0 - (\theta + c_3) y_1 \tag{S20}$$

$$y_3 = \kappa_2(x_2 y_0 + 2x_1 y_1 + x_0 y_2) - c_3\sigma^2 y_0 + c_3\sigma y_1 + c_3\sigma y_1 - (\theta + c_3) y_2 \tag{S21}$$

From Eq. (S19), we can find an expression for the CAR T-cell proliferation/exhaustion, $\kappa_2$, as,

$$\kappa_2 = \frac{y_1}{x_0 y_0} + \frac{(\theta + c_3)}{x_0} \tag{S22}$$

From Eq. (S20) and the expression for $\kappa_2$ in Eq. (S22), we can find an expression for $(\theta + c_3)$ as,

$$(\theta + c_3) = \frac{x_0 y_2}{x_1 y_0} - \frac{y_1}{y_0} - \frac{x_0 y_1^2}{x_1 y_0^2} - \frac{c_3 \sigma x_0}{x_1} \tag{S23}$$

From Eq. (S21) and the expression for $\kappa_2$ in Eq. (S22), we can find a complete expression for the effect of DEX on the CAR T-cell death rate, $c_3$, that no longer depends on any other unknown parameters. This expression is given as,

$$c_3 = \left[\frac{x_1}{\sigma x_2 y_0 + \sigma^2 x_1 y_0}\right] \left\{\frac{x_2 y_2}{x_1} + \frac{3y_1 y_2}{y_0} - \frac{x_2 y_1^2}{x_1 y_0} - \frac{2x_1 y_1^3}{x_1 y_0^2} - y_3\right\} \tag{S24}$$

Substituting the expression for $c_3$ from Eq. (S24) into Eq. (S23) results in a complete expression for the CAR T-cell death rate, $\theta$. Furthermore, substituting the expression for $(\theta + c_3)$ from Eq. (S23) and the expression for $c_3$ from Eq. (S24) into Eq. (S22) results in a complete expression for the CAR T-cell proliferation/exhaustion. Thus, all model parameters are identifiable. We note however that although the model parameters are identifiable, this does not mean the task of identifying them is straight forward. This is due to the extent to which the parameters $\kappa_2, \theta$, and $c_3$ exhibit nonlinear dependence on higher order derivatives of the functions describing the tumor cell and CAR T-cell populations. To perform parameter inference we employed a combination of Particle Swarm Optimization, the Levenberg-Marquardt algorithm, and iterative fitting procedures, each of which are summarized next.

### 7 Parameter estimation by particle swarm optimization

To estimate the model parameters from experimental data, we used a combination of particle swarm optimization (PSO) and the Levenberg-Marquardt algorithm (LMA). These optimization procedures were used to minimize the weighted sum-of-squares error between measured and predicted tumor cell and CAR T-cell populations. The objective function to be minimized is given as,

$$WSqE = \sum_{i=1}^{N} w_{i,x}[\tilde{x}_i - x_i(\xi)]^2 + w_{1,y}[\tilde{y}_1 - y_1(\xi)]^2 + w_{N,y}[\tilde{y}_N - y_N(\xi)]^2 \tag{S25}$$

where subscript $i$ represents the measurement time points, the $w_{i,x}$ and $w_{i,y}$ are the weights, the $\tilde{x}_i$ and $\tilde{y}_i$ are the measured tumor cell and CAR T-cell populations, and the $x_i(\xi)$ and $y_i(\xi)$ are the predicted tumor cell and CAR T-cell populations that vary based on the values of the model parameters $\xi = \{\rho, K, \kappa_1, \kappa_2, \theta, c_0, c_3\}$. Eq. (S25) has been written out to explicitly reflect the fact that measurements for the CAR T-cell population only exist for the initial and final time points. As the model Eqs. (5)-(6) are solved as an initial value problem, then to emphasize the importance of accuracy in the predicted final population of the CAR T-cell population, the corresponding weight for this term was set to $w_{N,y} = 10$, while all other weights were left with values of unity.

PSO is a stochastic global optimization procedure inspired by biological swarming [4]. PSO has been used recently for parameter estimation in a variety of initial value problems across cancer research and systems biology [5–8]. PSO treats the search for the global optimum of WSqE in Eq. (S25) as a moving swarm of fictitious particles, where each particle's coordinates represent values for the parameter set $\xi$. A weighted combination of the average swarm kinematics and the individual's historical kinematics are used to update the position of each particle. The update to the kinematics, specifically the position and velocity, are written explicitly as,

$$v_{i,j}(k+1) = \omega v_{i,j} + \phi_I r_I \left[\xi^I_{i,j}(k) - \xi_{i,j}(k)\right] + \phi_S r_S \left[\xi^S_{i,j}(k) - \xi_{i,j}(k)\right] \tag{S26}$$

$$\xi_{i,j}(k+1) = \xi_{i,j}(k) + v_{i,j}(k) \tag{S27}$$

where the subscripts $i$ and $j$ represent the particle number and parameter index, respectively, and $k$ represents the iteration step. The parameter values $\xi^I_{i,j}(k)$ and $\xi^S_{i,j}(k)$ are the parameter values for the individual and the swarm, respectively, that result in the best optimization of Eq. (S25). The constants $r_I$ and $r_S$ are random numbers between $[0, 1]$ to stochastically weight the influences of the individual best and the swarm best. The user can toggle the values of the three inputs $\omega, \phi_I$, and $\phi_S$. These inputs represent constant weights for inertial motion of an individual particle, and weights of the individual's and swarm's best historical parameter values. We implemented PSO with the python package pyswarm, using 50 particles, $\omega = 0.7, \phi_I = 2, \phi_S = 1$, a minimum parameter value step size of $1 \times 10^{-8}$, and terminated the procedure if either convergence was met within a tolerance of $1 \times 10^{-8}$, or 1000 iterations were completed [7, 8]. Once a global optimum of the model parameters was identified using PSO, a standard LMA was performed for finer resolution of the local optimization.

We employed an iterative fitting procedure to leverage the experimental control trials and reduce excessive guessing of upper and lower bounds on PSO parameters. For each tumor cell line, the tumor growth rate, $\rho$, and carrying capacity, $K$, are extracted from the untreated controls (neither DEX nor CAR T-cell treatment), using PSO-LMA. These values were then held constant for the PSO while determining the tumor killing rate, $\kappa_1$, CAR T-cell proliferation rate, $\kappa_2$, and CAR T-cell death rate $\theta$ for each initial CAR T-cell concentration untreated with DEX. The LMA step was then performed twice, once with the tumor cell growth rate and carrying capacity being held constant again, and once while letting them vary. The best resulting fit was then saved for analysis. A similar procedure was then used for each of the DEX concentration treatments.

2. Chiu CH, Lei KF, Yeh WL, Chen P, Chan YS, Hsu KY, et al. Comparison between xCELLigence biosensor technology and conventional cell culture system

for real-time monitoring human tenocytes proliferation and drugs cytotoxicity screening. Journal of orthopaedic surgery and research. 2017;12(1):1–13.
