## Supplementary figures and images for "Dose-dependent thresholds of dexamethasone destabilize CAR T-cell treatment efficacy"

### S1 Fig

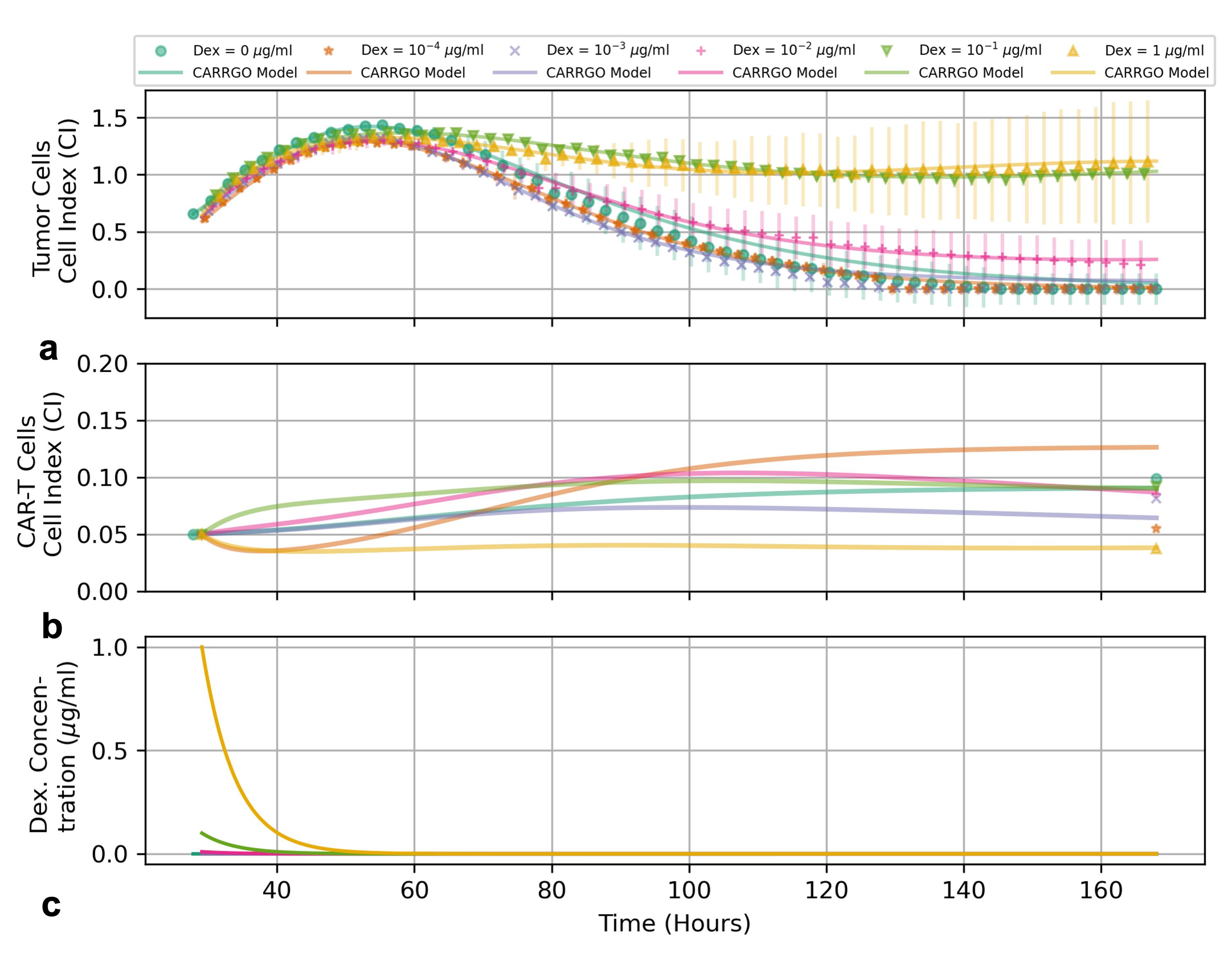

### S2 Fig

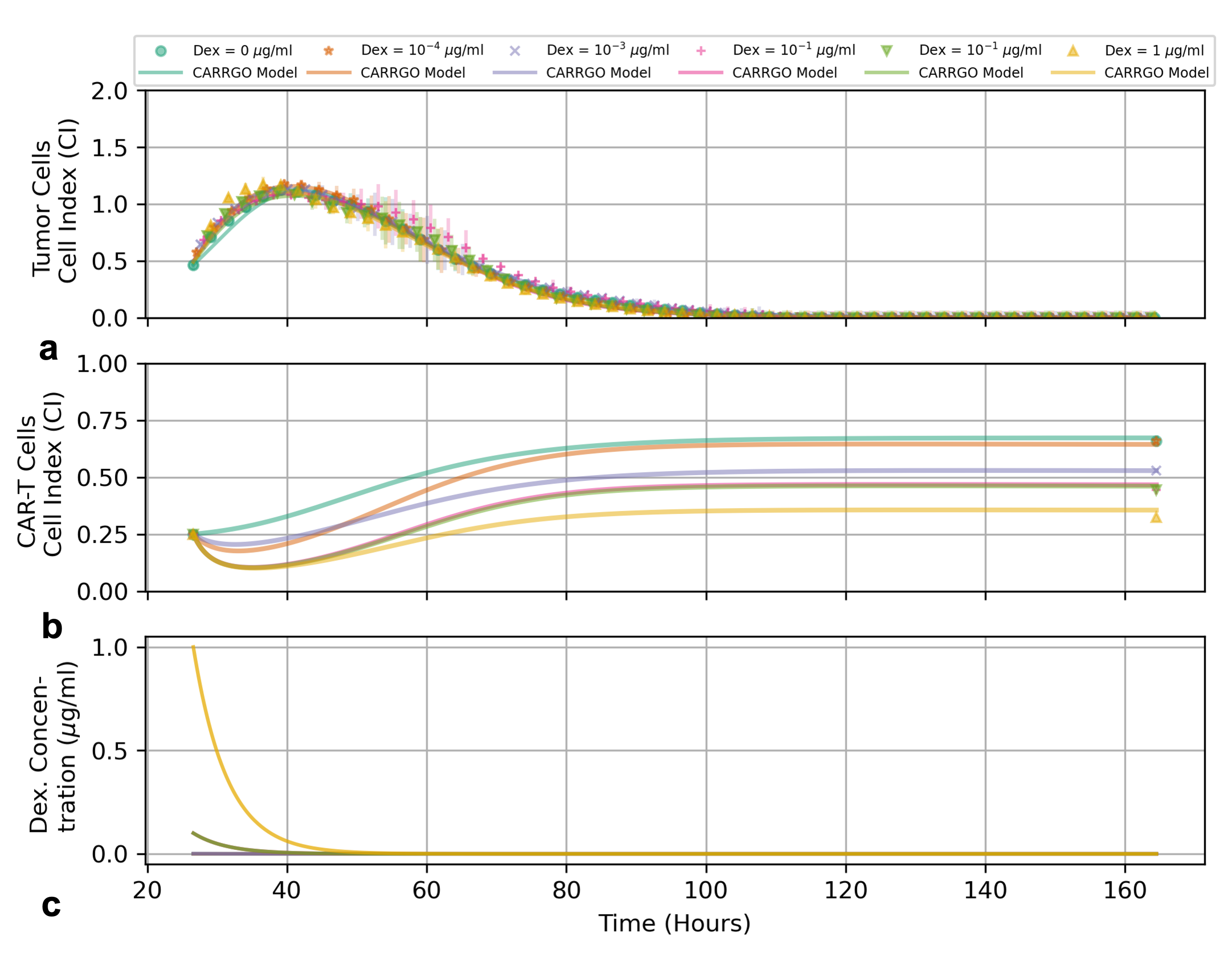

### S3 Fig

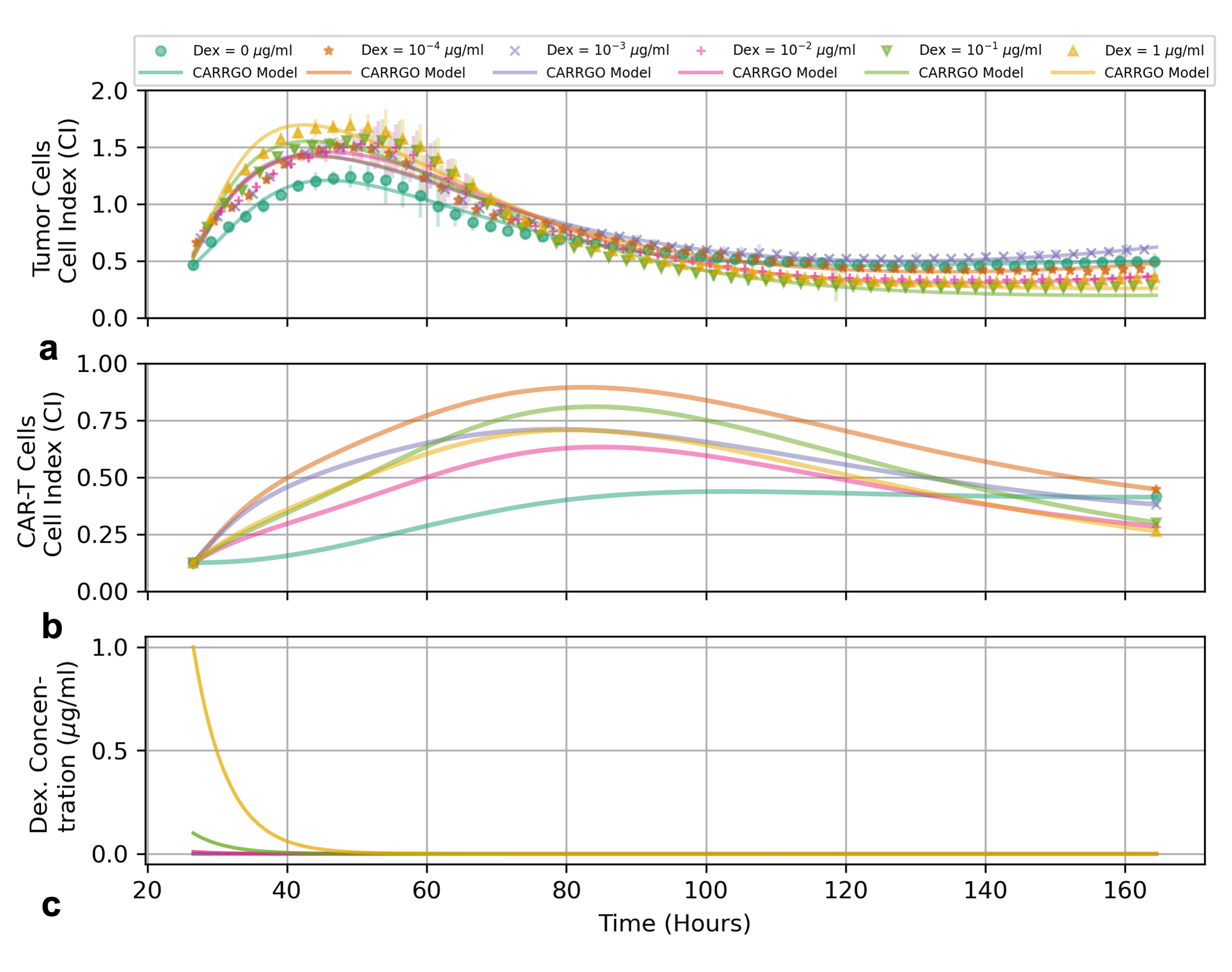

### S4 Fig

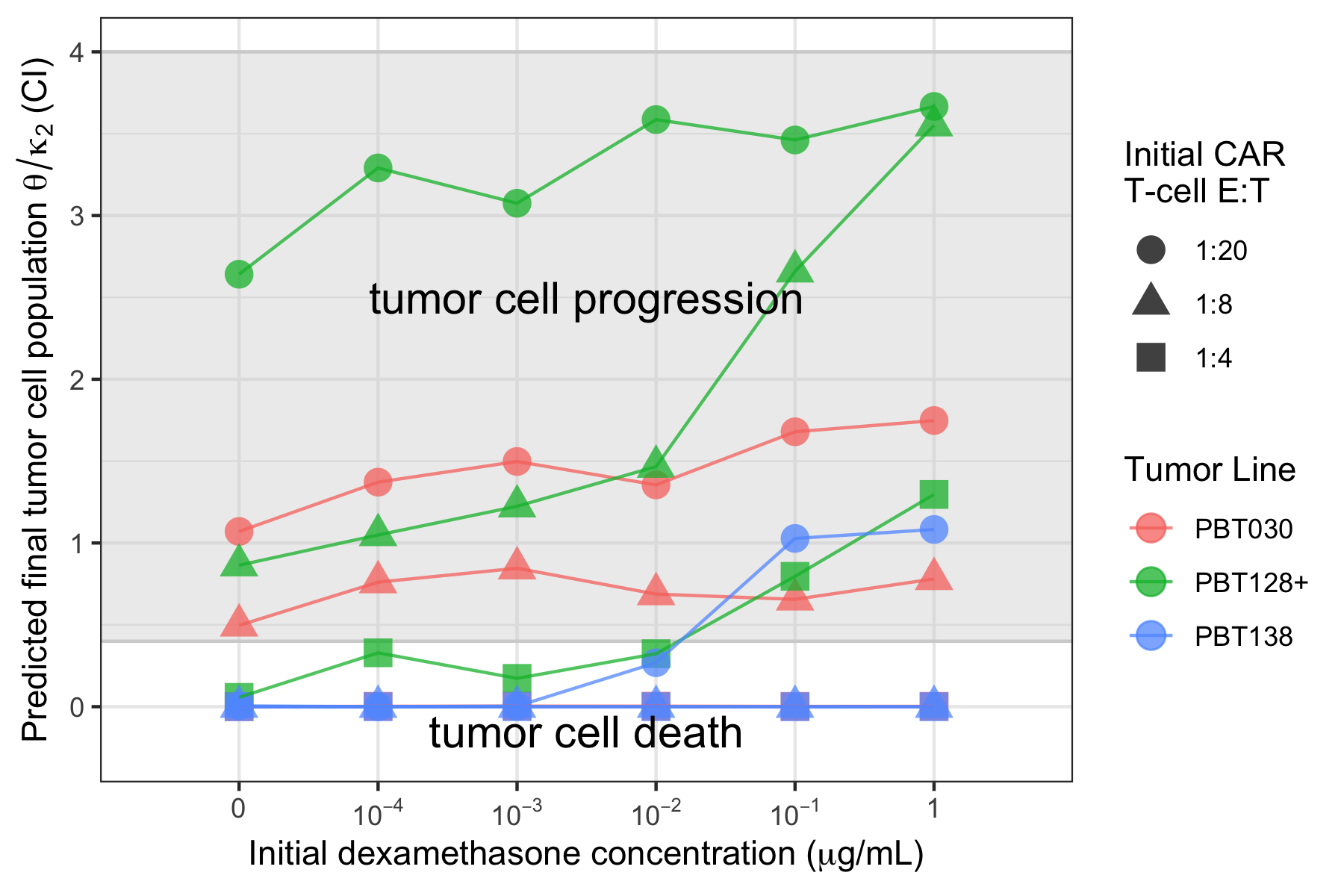
